# Macroscale traveling waves link perception, response selection, and vocal production during marmoset vocal interactions

**DOI:** 10.64898/2026.05.15.725341

**Authors:** Diya Yi, Xinyi Gao, Ruichen Tao, Misako Komatsu, Joji Tsunada

**Author notes:** **Co-corresponding authors:** Joji Tsunada, Chinese Institute for Bran Research, Beijing, C108, Bldg.3, No.9, YIKE Rd, Science Park Road, Zhongguancun Life Science Park, Changping District, Beijing 102206, China, Misako Komatsu, Institute of Science Tokyo, Tokyo, Japan.

## Abstract

Vocal communication involves a series of cognitive processes, which can be broadly categorized into three components: perceiving communicative signals; deciding whether and how to respond; and generating vocal motor output. These processes must work harmoniously, with integration and bridging between components being crucial for effective communication. Previous research on vocal communication has typically focused on specific brain regions or isolated cognitive functions, often lacking a holistic perspective of macro-scale, whole-cortical dynamics and their role in the complete communication process. Therefore, although the cortical areas associated with each cognitive component have been localized in humans, the macro-scale cortical dynamics underlying the integration of these cognitive processes remain unknown. Building on recent findings linking macro-scale cortical dynamics to behavioral performance, we hypothesized that traveling wave like cross-areal interactions play a role in integrating the three communicative components. To test this hypothesis, we recorded whole-cortical activity using epidural electrocorticography (ECoG) while subject marmosets vocally interacted with partners. We found theta-band activation in several cortical areas, including the parietal and auditory cortices, while listening to partner’s calls. This activity was further modulated depending on whether the subjects engaged in vocal interactions, potentially representing the transformation of sensory processing into decision-making and vocal motor preparation. Given the widespread nature of this modulation, we next characterized whole-brain activity patterns by employing a novel analytical method, Weakly Orthogonal Conjugate Contrast Analysis (WOCCA). This analysis revealed that cortical activity could be decomposed into two distinct traveling wave like propagation patterns, a rotational and a translational wave, and both waves discriminated communicative conditions consistent with localized activity. The rotational wave further represented vocal motor preparation through trigger-like temporal pattern. In addition, the magnitude of the translational wave immediately before subject’s vocal production correlated with the vocal production-induced suppression of high-gamma-band activity, particularly in the prefrontal and auditory cortices. As vocalization-induced suppression is believed to reflect sensory prediction, the translational wave may propagate specific decision-related or acoustic information necessary for subsequent vocal production to local cortical areas. These findings suggest that the brain orchestrates the sequential cognitive processes underlying vocal communication through macro-scale traveling waves.

## Introduction

Vocal communication relies on a sequence of cognitive processes that can be broadly grouped into three components: (1) perceiving communicative signals, requiring the separation, recognition, and categorization of speech to comprehend meaning; (2) deciding whether and how to respond, involving decision-making and vocal planning; and (3) generating vocal motor output, which includes initiating and coordinating speech production^1–6^. For effective communication, these components must be coordinated, with integration and bridging between them ensuring smooth transitions from perception to decision to action^4,5,7–13^. While prior research has identified cortical areas associated with each component in humans and non-human primates, a holistic perspective on macro-scale, whole-cortical dynamics—and how they integrate these processes across distributed networks—remains limited^4,6,14–53^.

Foundational work in human neuroscience localized key cortical substrates of speech perception and production (e.g., classical Broca and Wernicke areas), and recent neurophysiological studies have refined this view by delineating pathways and temporally structured activations across superior temporal, frontal, and motor cortical regions during speech perception, planning, and execution^4,14–18,20,26–39^. However, these studies largely emphasize region-specific activity or isolated computations. The macro-scale cortical dynamics that integrate sensory processing with decision-making and vocal motor preparation, particularly their cross-areal interactions during naturalistic social vocal exchanges, are still poorly understood.

Marmoset monkeys (*Callithrix jacchus*) provide a powerful model for investigating the cortical mechanisms of vocal communication at scale^6,54–56^. Marmosets exhibit human-like aspects of vocal control, including contingent timing of responses and compensation for environmental interference, enabling controlled experiments that link behavior to neurophysiology^57–62^. Prior work has implicated auditory cortex in processing conspecific vocalizations and monitoring auditory feedback, and frontal areas in generating vocal motor commands^6,25,41–45,48,54,63–71^. Yet, a core question remains: how does the cortex integrate these distributed functions across time and space to orchestrate the sequential processes of vocal communication?

To address this, we recorded whole-cortical activity using epidural electrocorticography (ECoG)^72,73^ while visually isolated marmosets engaged in natural vocal interactions with partners. We first identified theta-band activation across multiple cortical regions, including auditory and parietal cortices, during listening to partner calls. Critically, this sensory activity was modulated by whether subjects engaged in vocal interactions, suggesting the transformation of sensory processing into decision-making and vocal motor preparation. Motivated by the widespread nature of these modulations and recent work linking macro-scale dynamics to behavior^74–76^, we hypothesized that traveling wave like cross-areal interactions play a role in integrating the three communicative components. To test this hypothesis, we applied a novel analytical method, Weakly Orthogonal Conjugate Contrast Analysis (WOCCA)^77^, to characterize whole-brain activity patterns. WOCCA revealed two distinct traveling wave–like propagation modes: a rotational pattern corresponding to vocal motor preparation and a translational pattern conveying sensory prediction signals associated with vocal production. Together, these findings suggest that macro-scale traveling waves link the sequential cognitive processes underlying vocal communication by integrating sensory, decision, and motor components across the cortex.

## Results

### Timing control of antiphonal calling is the core of vocal interactions under visual isolation

We recorded natural antiphonal exchanges under visually isolated conditions and detected 6904 calls from subject Wa and its partner and 4196 calls from subject Sa and its partner (Fig. 1a). During these interactions, marmosets predominantly produced phee calls (Wa: 1371; Wa partner: 3596; Sa: 759; Sa partner: 2078), and subsequent analyses focused on this call type.

**Figure 1.**
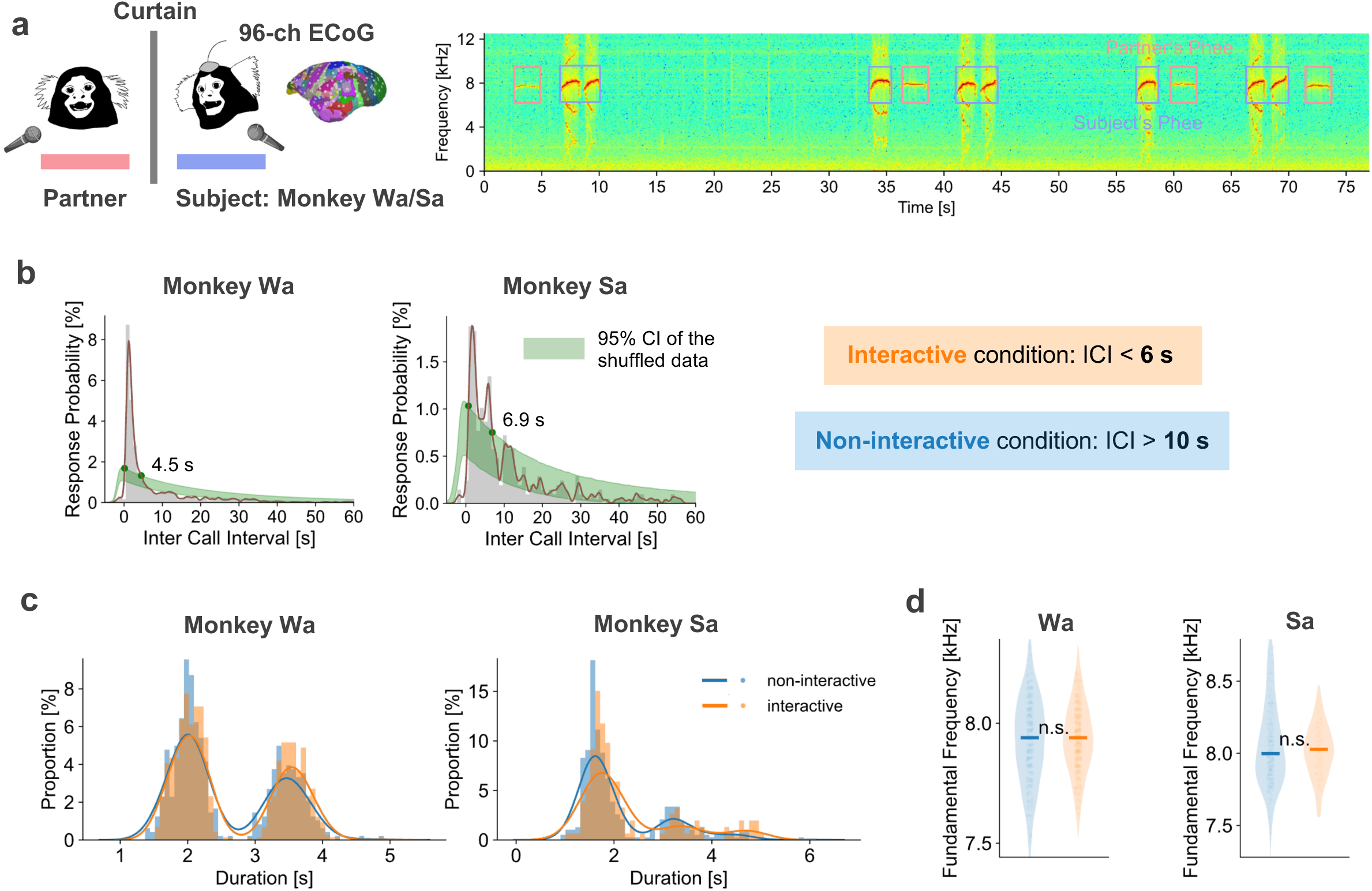
Behavioral characterization of antiphonal vocal interactions. (a) Experimental setup and example sound spectrogram illustrating calls produced by an ECoG-implanted subject and an unimplanted partner (ECoG schematic inset). Monkeys were visually isolated and separated by 2.5 m. (b) Inter-call-interval (ICI) distributions for call sequences from partner → subject. Left: Monkey Wa; right: Monkey Sa. Green shading denotes the 95% confidence interval from temporally shuffled data; green dots indicate ICI bins where the empirical distribution exceeded the shuffle distribution. Based on these distributions, we defined interactive responses as ICI < 6 s and non-interactive calls as ICI > 10 s. (c,d) Call structure and acoustics did not differ between interactive and non-interactive contexts. (c) Call duration distributions for Wa (left) and Sa (right), separated by one-phrase and two-phrase calls (Wilcoxon rank-sum test; for Wa, p=0.541 for the one phrase calls and p = 0.081 for two phrase calls and for Sa, p<0.01 for the one phrase calls and p=0.978 for two phrase calls). (d) Fundamental frequency distributions for Wa and Sa (Wilcoxon rank-sum test, p=0.778 for Wa, p=0.809 for Sa).

Consistent with prior reports^57,58^, a hallmark of antiphonal calling is a peak in the inter-call interval (ICI) distribution for successive calls produced by different individuals (e.g., partner → subject) within approximately 10 s. We computed ICIs between partners’ calls and the subjects’ subsequent calls, and normalized the ICI distributions by the number of partner calls to estimate response probability as a function of latency (Fig. 1b). For both subjects, observed ICI distributions exceeded time-shuffled controls for ICIs (ICI < 4.5 s for monkey Wa, < 6.9 s for monkey Sa, bootstrap test, p < 0.05 for both), with a prominent peak shortly after partners’ call offset (1.2 s for Wa, 1.7 s for Sa). Based on these distributions, we classified calls with ICI < 6 s as interactive. To conservatively define non-interactive calls, we used ICI > 10 s. Interactive-response probabilities to preceding partner’s calls by subjects were: 20.5% for Wa and 7.4% for Sa. Although there were individual differences (e.g., Sa’s overall response probability was lower than Wa), these differences likely reflect subject–partner pairing and subjects’ spontaneous call rates.

To estimate spontaneous production tendencies independent of partner timing, we analyzed within-subject ICIs for repetitive call onsets (subject → subject). These distributions differed from cross-individual ICIs, exhibiting a later, broader peak consistent with slower, intrinsic rhythms potentially related to cardiorespiratory/Mayer-wave influences^78–80^.

We next asked whether acoustic structure differed between interactive and non-interactive calls. We found no significant differences in duration (Fig. 1c) or fundamental frequency (Fig. 1d). These results indicate that, in our paradigm, antiphonal calling is primarily defined by precise timing control rather than changes in call acoustics, providing a behavioral substrate for subsequent analyses of decision and motor preparation.

### Different macroscale cortical activity patterns during sensory processing of partners’ vocalizations in interactive vs. non-interactive contexts

We analyzed whole-cortical ECoG during antiphonal calling. Due to wireless recording constraints, call numbers used for each epochs varied as shown in Table S1, but at least, neural analyses included 1140 calls from Wa, 647 from Sa, and 4652 (Wa partner 2836, Sa partner 1816) partner calls. Time–frequency analyses aligned to partner vocalizations revealed robust, distributed sensory responses spanning auditory, parietal, and prefrontal cortices in both subjects (Fig. 2a). Those sites showed phasic increases in theta (5-8 Hz) power, while parietal sites showed sustained theta increases. On the other hand, high-gamma (50-150 Hz) power increased in auditory sites, but suppressed in parietal and prefrontal sites.

**Figure 2.**
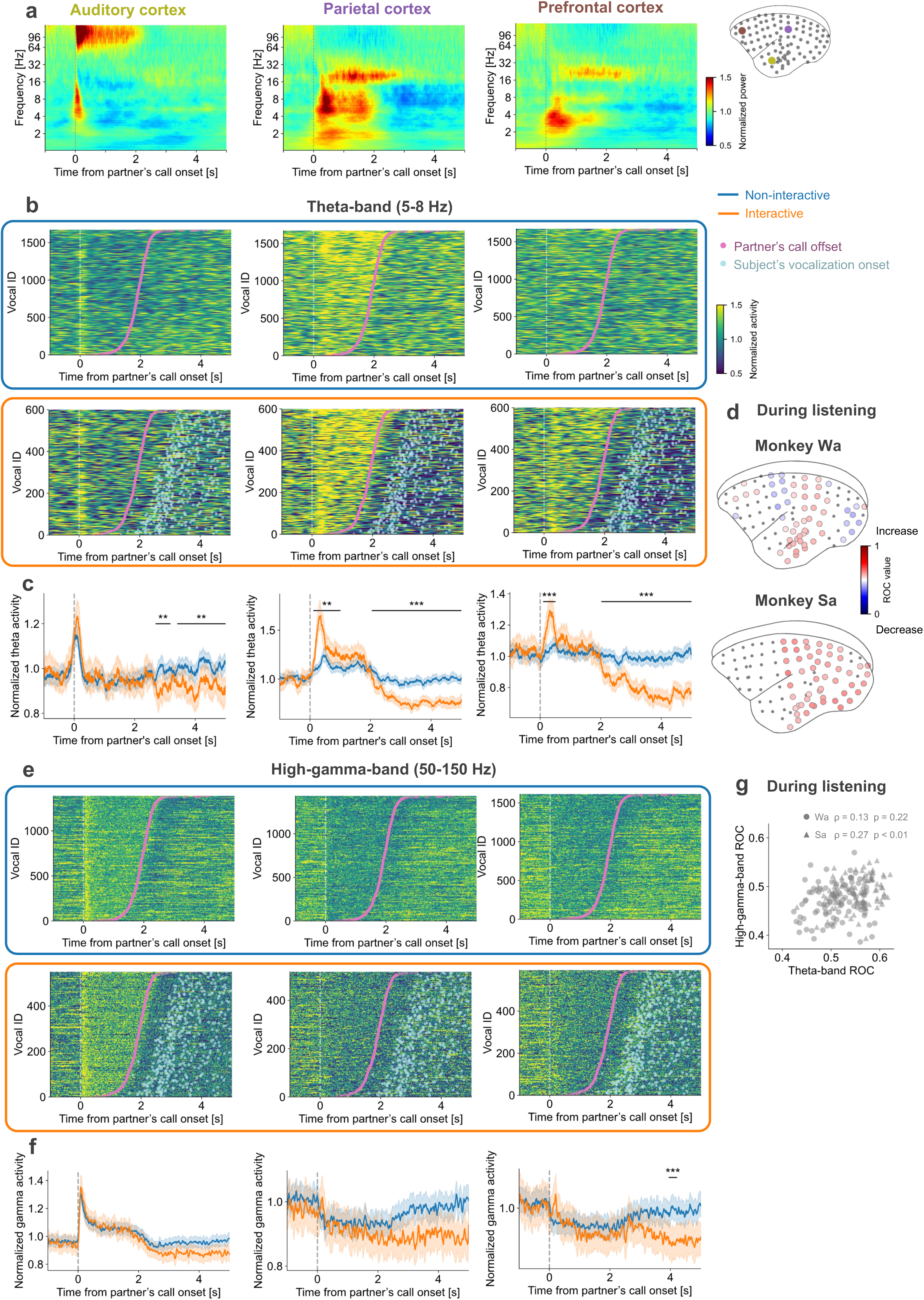
Sensory responses to partners’ vocalizations are modulated by whether the subject respond to the partner. (a) Time-frequency plots of mean ECoG power recorded from representative auditory, parietal, and prefrontal electrodes in monkey Wa, aligned to partner’s call onset. Data are pooled across sessions (partner calls: N=2836 for Wa, N=1816 for Sa). Electrode locations are indicated on the cortical schematic (inset). (b,c) Trial-level theta-band activity and averages during partner’s calls for non-interactive and interactive contexts. (b) Each row shows one call (sorted by call duration). Pink markers indicate partner call offset; light-blue markers indicate onset of the subject’s response call (when present). Top (bottom) panels: non-interactive (interactive) context. (c) Median normalized theta-band activity for non-interactive (blue) and interactive (orange) conditions; shading denotes 95% confidence intervals (bootstrap, n=1000). Asterisks mark time points/windows with significant differences between conditions (Wilcoxon rank-sum test, * p<0.05, ** p<0.01, *** p<0.001). (d) Whole-cortex maps of context preference during partner-call epochs quantified as ROC values comparing interactive vs. non-interactive trials (top: Wa; bottom: Sa). ROC values greater than 0.5 (less than 0.5) indicate higher activity in the interactive (non-interactive) condition. Only electrodes with significant effects are shown (permutation test, p<0.05). (e,f) High-gamma-band activity shown in the same format as (b,c). (g) Relationship between electrode-wise context preference (ROC values) in theta and high-gamma bands. A significant cross-band relationship was observed in monkey Sa (Spearman’s rank correlation, rho ρ=0.27, p<0.01), whereas Wa showed no reliable correlation (rho ρ=0.13, p=0.22). Across animals, interactive trials tended to show stronger theta-band responses (ROC > 0.5) and weaker high-gamma responses (ROC < 0.5, Wilcoxon rank-sum test, p<0.05 for both monkeys and for both frequency bands).

Critically, sensory-evoked activity depended on communicative context, defined behaviorally by response timing (interactive: subject responded within 6 s of the partner call; non-interactive: no response within 10 s). Across the cortical surface, theta-band responses were significantly larger in interactive than non-interactive contexts (Fig. 2b,c,d). In contrast, high-gamma modulations were less frequent and spatially sparser (Fig. 2e,f). The magnitudes of theta- and high-gamma-band contextual modulations were not consistently correlated across sites (Fig. 2g). This may be partially explained by complex modulation pattern by the combination of cortical sites and frequency bands: overall increase in theta-band activity during interactive context across broad recording sites vs. mixture of sensory-evoked excitatory (e.g., auditory cortex) and suppressive responses (e.g., prefrontal and parietal cortices) with further contextual modulations in the high-gamma band. Together, these findings indicate that widespread theta-band activity is selectively enhanced when subjects are likely to engage in vocal interaction, whereas high-gamma changes are more focal, suggesting different roles for low- and high-frequency activity in linking sensory processing to subsequent decision and motor preparation.

### Macroscale cortical activity patterns related to vocal production in interactive and non-interactive contexts

Neural activity around subjects’ own vocal production differed markedly from sensory-evoked responses to partners’ calls (Fig. 3a). In the pre-vocal period (−1 to 0 s relative to vocal onset), high-gamma power exhibited a complex profile, with a slower suppression superimposed on a tonic increase, most prominently in prefrontal and premotor regions, whereas theta power decreased. In contrast, the auditory cortex showed pre-vocal suppression of high-gamma activity as reported before^48,70,71^. During vocalization, both theta and high-gamma activity were broadly suppressed across auditory, parietal, and prefrontal cortices, consistent with a global change in cortical state during self-generated vocal output^48,66,70,71,81^. We additionally observed beta-band increases at a subset of sites in auditory and parietal cortex, but this effect was evident only in monkey Wa.

**Figure 3.**
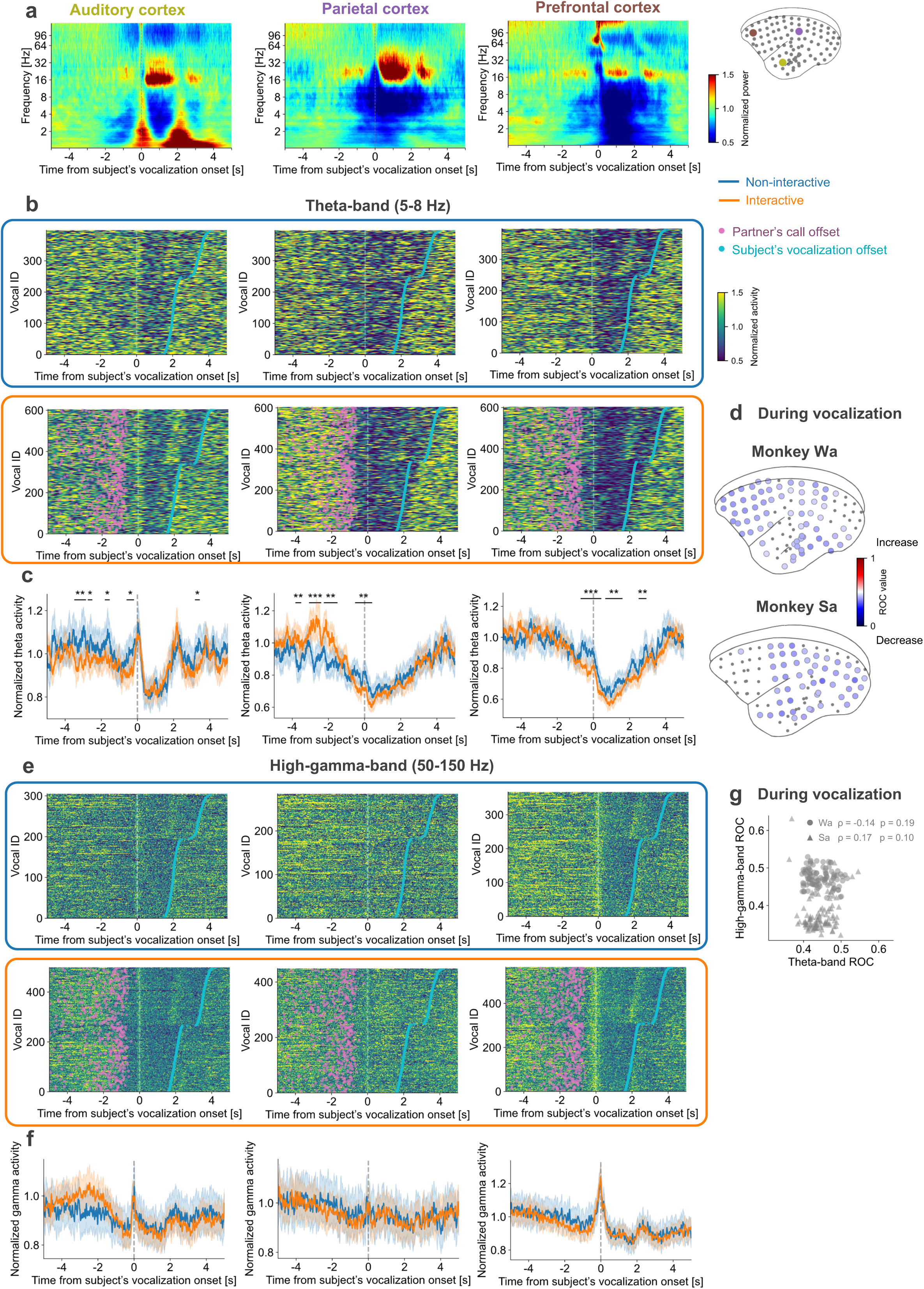
Neural activity around subject vocal production is modulated by communicative context. (a) Time–frequency plots of mean ECoG power from representative auditory, parietal, and prefrontal electrodes in monkey Wa aligned to subject call onset. Data are pooled across sessions (subject calls: N=1140 for Wa, N=647 for Sa). Electrode locations are indicated on the cortical schematic (inset). (b,c) Trial-level theta-band activity and averages for non-interactive and interactive subject calls. (b) Pink markers indicate the offset of the partner’s preceding call (when present); cyan markers indicate the subject call offset. Top (bottom) panels: non-interactive (interactive) context. (c) Median normalized theta-band activity for non-interactive (blue) and interactive (orange) conditions; shading denotes standard errors. Asterisks indicate significant differences (Wilcoxon rank-sum test, * p<0.05, ** p<0.01, *** p<0.001). (d) Whole-cortex maps of context preference (ROC values) during vocal production. ROC values greater than 0.5 (less than 0.5) indicate higher activity in the interactive (non-interactive) condition. Only significant electrodes are displayed (permutation test, p<0.05). Consistent with these three electrode examples, global suppression, with additional suppression for interactive context, was observed in the theta-band. (e,f) High-gamma-band activity shown in the same format as (b,c). (g) Relationship between electrode-wise context preference in theta and high-gamma bands during vocalization. No reliable cross-band correlation was observed, but both bands exhibited stronger suppression in the interactive context during pre-vocal and vocalization periods (Wilcoxon signed-rank test, p<0.05).

Vocal production-related activity was also modulated by communicative context. We defined interactive productions as subject calls emitted within 6 s of a partner’s preceding call, and non-interactive productions as calls with ICI > 10 s, which were presumably spontaneous. Comparing these contexts first revealed that theta-band suppression before and during vocalization was stronger and more widespread for interactive than non-interactive productions (Fig. 3b,c,d), extending across broad cortical areas. Second, pre-vocal increase in prefrontal high-gamma band did not show contextual modulations (Fig. 3 e,f), but overall suppression during vocalization was stronger and broadly observed for interactive context (Fig. 3e,f,g).

Finally, to test whether theta and high-gamma modulations reflect a shared control process, we assessed the across-site relationship. This analysis revealed, in spite of the overall suppression for both theta and high-gamma bands in interactive production, no significant correlations during vocal period (Fig. 3g), suggesting that the two bands capture independent aspects of the context dependency during vocal production.

Together with the context-dependent theta modulation observed during listening to partner’s calls, these pre-vocal and vocal patterns suggest that interactive exchanges engage a distributed preparatory state, expressed as widespread theta and high-gamma suppression, linking communicative context to upcoming vocal motor output.

### Differential activity related to vocal interactions forms macro-scale traveling wave dynamics

The context-dependent modulations observed during listening (Fig. 2) and vocal production (Fig. 3) were distributed across broad cortical areas, raising the possibility that vocal interactions engage coordinated, macro-scale dynamics that integrate sensory, decision, and motor processes. As an initial test of large-scale coordination, we quantified the functional connectivity across cortical sites using mean activity during each behavioral epoch. During sensory processing of partner calls, theta-band activity exhibited structured positive correlations across multiple cortical areas, forming clusters of sites with coherent response profiles (Fig. S1). Interestingly, these correlation structures did not change significantly even during the pre-vocal and vocal periods. Furthermore, comparing interactive and non-interactive contexts revealed no significant differences (Fig. S1), indicating that communicative context alters local response amplitude but not the population-level coordination across the cortex or mean-activity based connectivity analysis may be not sensitive to capture the temporally dynamic contextual modulations as discussed next.

Although our correlation-based functional connectivity analysis provided evidence of the coherent ECoG activity across cortical areas, this analysis does not resolve the directional and temporally fine-grained propagation expected for information integration across distributed networks. Therefore, we strategically decided to use the traveling wave analysis, enabling us to extract complex activity propagation patterns on call-by-call basis with fine temporal resolution^77^.

This approach was necessary because cortical activity during a partner’s call was highly dynamic (Fig. 4a) and could not be adequately summarized by static, region-averaged responses. For example, parietal and auditory electrodes, showing strong sensory responses (Fig. 2), exhibited a reproducible theta-band phase gradient that progressed from parietal cortex toward auditory cortex via middle temporal regions, consistent with traveling-wave–like propagation. This method also fit well with our purpose of characterizing multi-site propagation motifs over the cortex, in comparison with several other functional connectivity methods (Granger causality, transfer entropy, and phase-slope index), typically giving insights into the pair-wise activity transmission. Briefly, here, the traveling wave is defined as a pattern of entire-cortical oscillations with a spatial phase gradient across cortical areas. The traveling wave analysis decomposes the entire-cortical oscillations into directionally independent phase gradient by using the weakly orthogonal conjugate contrast analytical method (WOCCA; Fig. 4b)^77^. Similar to the principal component analysis (PCA), this analysis gives us traveling wave components with explained traveling energy (similar as explained variance in PCA), allowing us to extract the pattern and contribution of each wave. In addition, WOCCA allows us to extract pure propagation patterns while minimizing spurious standing-wave components that can arise in traditional methods. For each component, the sign of the projection value defines the dominant direction, while its magnitude quantifies the strength of that wave pattern.

**Figure 4.**
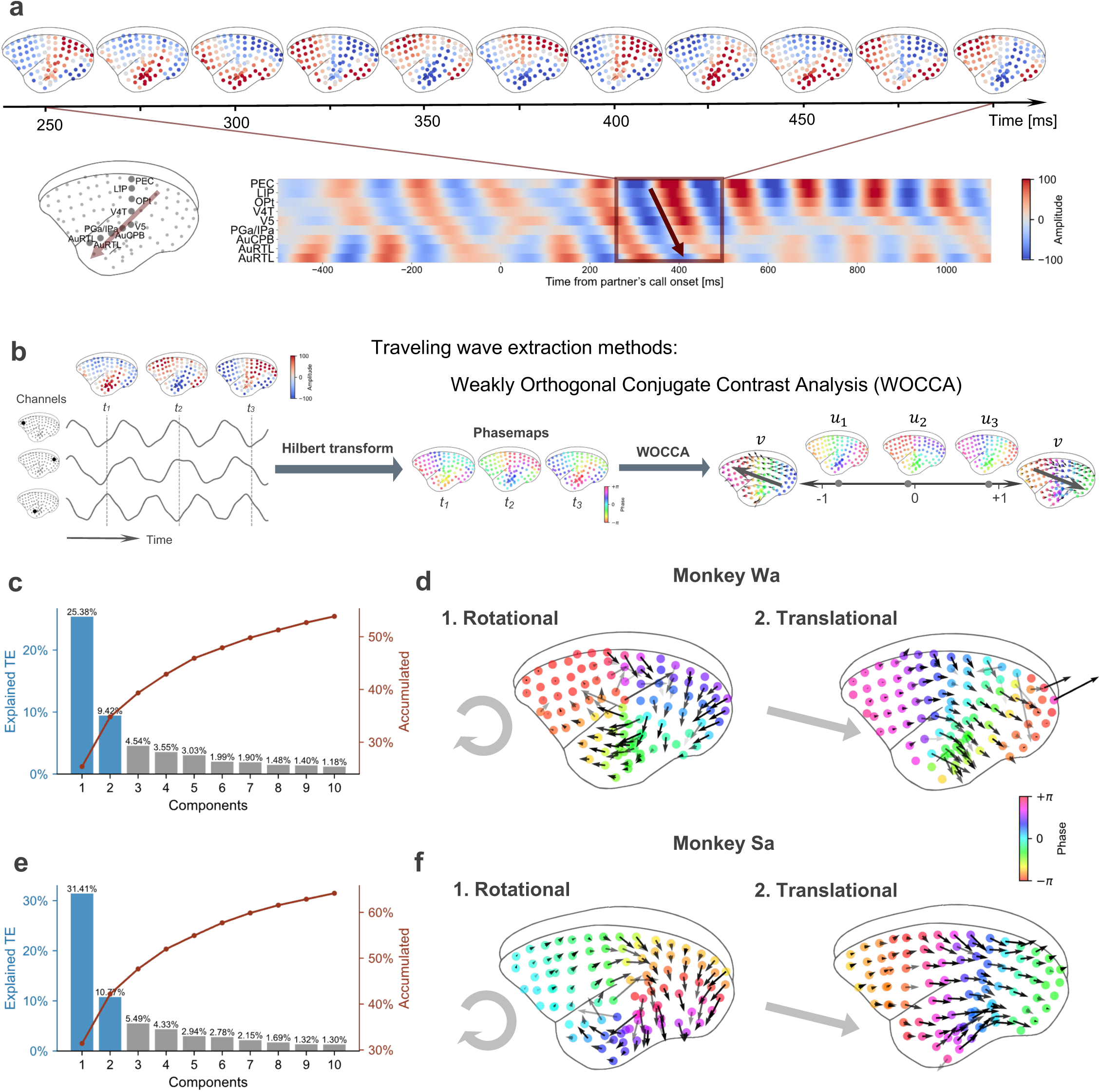
Whole-cortex oscillations exhibit traveling-wave structure captured by WOCCA. (a) Example theta-band activity across electrodes during a partner call illustrating a spatiotemporally organized phase gradient consistent with traveling-wave propagation. Colors show band-limited theta activity. (b) Overview of the traveling-wave analysis using Weakly Orthogonal Conjugate Contrast Analysis (WOCCA)^77^. WOCCA decomposes whole-cortex oscillatory activity into traveling-wave components characterized by spatial phase-gradient templates and trial-resolved projection values. (c) Explained traveling-energy for each component (blue bars) and cumulative explained traveling energy (red line) for each monkey (Wa, top; Sa, bottom). The first two components accounted for the largest proportion of variance (Wa: 35%, Sa: 42%), and were used for subsequent analyses. (d) Spatial phase-gradient templates for components 1 and 2 for Wa (top) and Sa (bottom). Colors indicate relative phase across electrodes. Arrows depict the local phase-gradient direction. Templates were conserved across monkeys after merging the electrodes layout (complex correlation coefficient [CCC]: magnitude of the complex inner product between normalized vectors, comp1 CCC=0.879, comp2 CCC=0.826, p<0.001 for both). Component 1 and component 2 were labeled as rotational and translational patterns, respectively, based on their spatial organization.

Applying WOCCA revealed two major traveling-wave components that together accounted for significant proportions of variance in the analyzed oscillatory activity (Wa: 35%; Sa: 42%; Fig. 4c,e). After merging the electrodes layout, the spatial phase-gradient patterns were consistent across subjects (complex correlation coefficient [CCC]: magnitude of the complex inner product between normalized vectors, comp1 CCC=0.879, comp2 CCC=0.826, p<0.001 for both), indicating reproducible macro-scale propagation modes.

Component 1 corresponded to a phase-gradient pattern consistent with propagation along a parietal → temporal → frontal pathway for positive projection values, and the opposite direction (frontal → temporal → parietal) for negative values (Fig. 4 d,f left). Component 2 expressed a distinct phase-gradient pattern consistent with a frontal → temporal/parietal pathway for positive projection values and the opposite direction for negative values (Fig. 4 d,f right).

How does each component change its wave amplitude and direction during vocal interactions? During listening to partner’s calls, component 1 showed larger positive projection values in the interactive context, with a relatively delayed peak (approximately 2s after call onset) and sustained elevation extending beyond call offset (Fig. 5a,c). This context dependence was present in both monkeys (p<0.05 for both monkeys). Notably, the interactive vs. non-interactive difference diminished during the pre-vocal and vocal epochs, although component 1 exhibited a clear pre-vocal build-up toward vocal onset (Fig. 5e).

**Figure 5.**
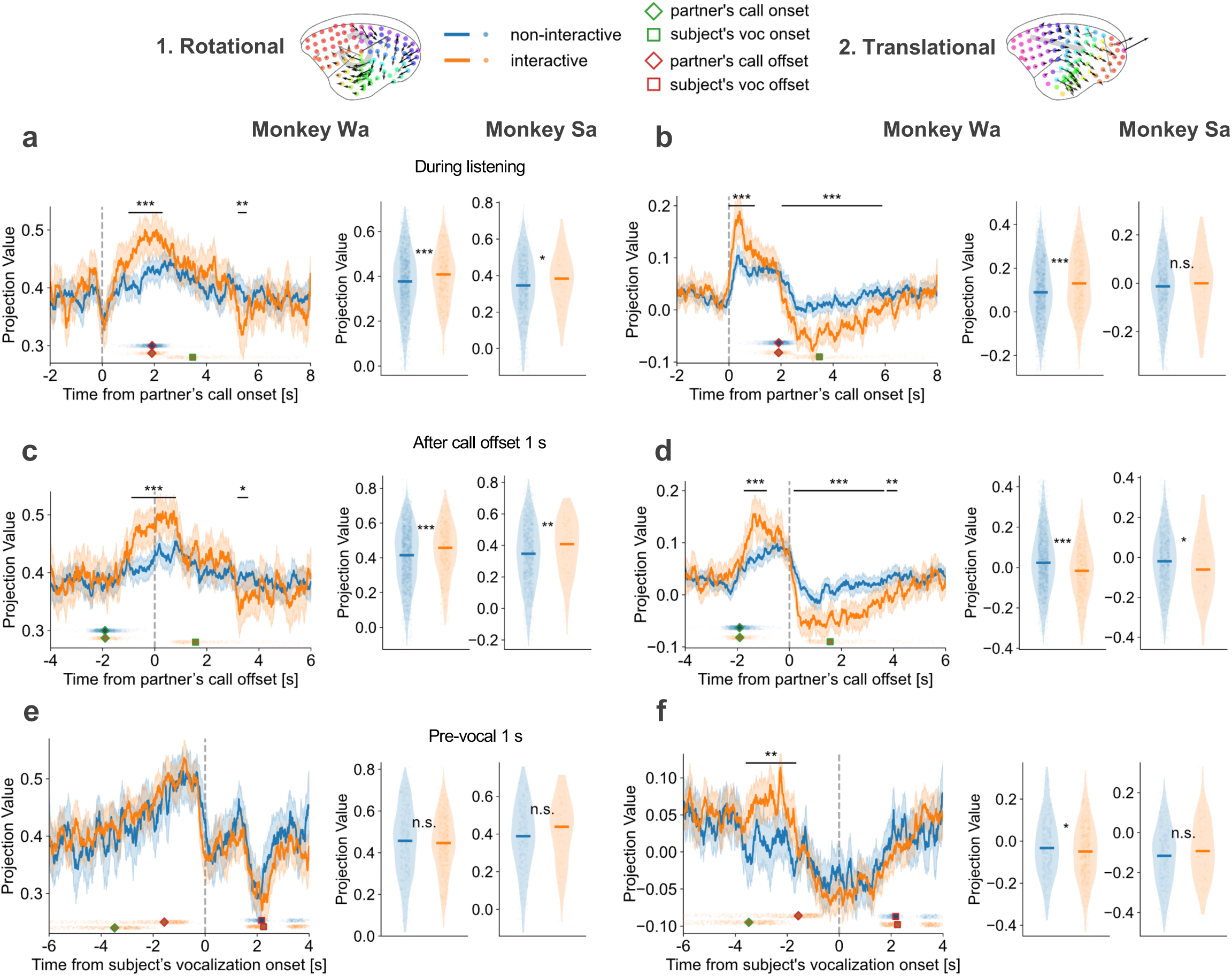
Traveling-wave components discriminate communicative context with distinct temporal profiles. (a,c,e) The time course and quantification of traveling wave component 1 (rotational wave) aligned to partner’s call onset (a), partner’s call offset (c), and subject’s call onset (e). Projection values are signed: the sign indicates whether the instantaneous phase gradient matches the template direction (positive) or the opposite direction (negative), and the magnitude reflects wave strength. Left subpanels: median (±95% confidence interval, bootstrap, n=1000) time courses for non-interactive (blue) and interactive (orange) trials. Asterisks mark significant differences between contexts (Wilcoxon rank-sum test, * p<0.05, ** p<0.01, *** p<0.001). Event-time distributions are overlaid as markers (partner’s call onset/offset; subject’s call onset/offset as labeled in each panel). Middle/right subpanels: window-averaged projection values for Wa and Sa, respectively. The analysis windows were call duration (a), the interval from partner’s call offset to one second after offset (b), and the one-second pre-vocal period (c). Asterisks indicate statistical significance (Wilcoxon rank-sum test, * p<0.05, ** p<0.01, *** p<0.001). (b,d,f) Results of traveling wave component 2 (translational wave) are shown in the same format as (a,c,e). Component 2 exhibits earlier context dependence during partner-call processing and a sign reversal around partner-call offset. It should be noted that, for the time course plots, some context differences necessarily co-vary with call availability (e.g., interactive trials contain a partner call followed by a subject response, whereas non-interactive trials may contain only a single animal’s call).

In contrast to component 1, component 2 exhibited earlier context dependence during partner’s calls, peaking around 0.5 s after partner call onset (Fig. 5b). Strikingly, the sign of component 2 projections reversed around partner call offset, suggesting a rapid switch in the dominant propagation direction at a behaviorally relevant transition point (from ongoing sensory analysis to response selection and/or preparation; Fig. 5d). During the pre-vocal and vocal epochs, component 2 projections were predominantly negative and showed little context dependence, consistent with a common production-related process engaged regardless of whether the call was interactive or spontaneous (Fig. 5f). Although component 2 expressed both signs over time, positive projections were larger in magnitude than negative projections during listening, supporting its primary role in early, context-dependent processing of partner calls (Fig. 5b). Moreover, together with its earlier peak relative to component 1, the propagation direction of component 2 (frontal to posterior cortices) is consistent with a role in rapid response selection/decision-related processing during partner’s calls in the interactive context (Fig. 5a,b).

Although both components showed prominent peri-vocal modulation (Fig. 5e,f), they differed qualitatively: component 1 exhibited a monotonic build-up toward vocal onset, whereas component 2 showed sign switching across listening and production epochs, supporting the interpretation that they capture distinct macro-scale modes rather than a single shared factor. These findings indicate that vocal interaction recruits at least two separable traveling-wave dynamics with different timing, directionality, and relationships to behavioral context.

### Traveling waves predict the timing of subject’s vocal responses and are associated with vocalization-induced suppression

To further evaluate the functional relevance of the traveling-wave components, we tested whether trial-by-trial wave projections predicted key features of subjects’ vocal behavior during interactive exchanges. We quantified relationships between component projection values and subsequent vocal output using correlation analyses. Specifically, we examined: 1) the peak timing of component 1 relative to subject’s vocal onset, and correlations between component 2 and 2) the inter-call interval (ICI; partner → subject), 3) the subject call’s fundamental frequency, and 4) call duration.

For component 1’s pre-vocal build-up, when trials were randomly selected independent from inter-call intervals, component 1’s projection values peaked around 1s before vocal production for both monkeys (786.8 ms for Wa and 1159.2 ms for Sa; bootstrap test, p<0.05; Fig. 6a), suggesting that this component reflects a preparatory process that builds toward a threshold before vocal initiation.

**Figure 6.**
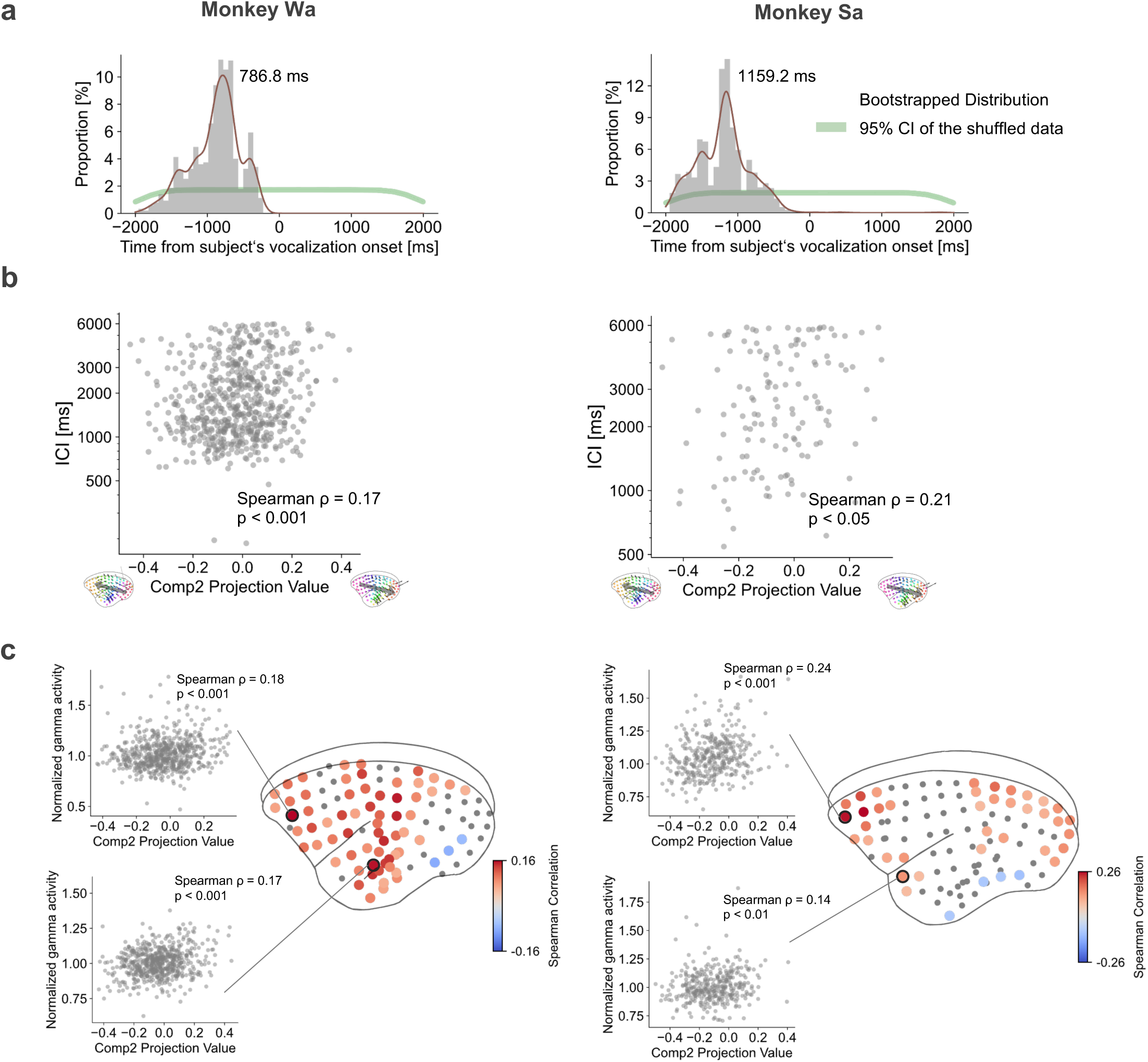
Traveling waves predict response timing and covary with vocalization-induced suppression. (a) Distributions of peak projection values of traveling wave component 1 relative to subject’s vocal onset for monkey Wa (left) and Sa (right). Peaks were calculated by repeatedly subsampling 120 subject calls (1500 iterations). Green shading indicates the 95% confidence interval obtained from temporally shuffled data (1000 shuffles). Peak projection values appeared ∼1 s before vocal onset, consistent with a preparatory build-up. (b) Component 2 projection values predict response timing. Projection values computed in the window from partner call offset to +1 s correlated with ICI for both monkeys (Spearman’s rank correlation, p<0.05). Negative vs. positive projections correspond to opposite propagation directions (see insets for the template patterns), and were associated with shorter vs. longer ICIs, respectively. This anlysis focused on interactive condition only (i.e., ICI < 6 s). (c) Electrode-wise correlations between pre-vocal high-gamma suppression and component 2 projection values shown as scatterplots and also mapped onto the cortical surface. Only significant electrodes are shown (Spearman correlation, p < 0.05). Significant relationships were prominent in auditory, parietal, and frontal regions, indicating coupling between a macro-scale traveling-wave component and vocal production-related local high-gamma suppression.

Furthermore, across monkeys, component 2 projection values after partner’s call offset (1s interval after call offset) showed reliable relationships with response timing (Fig. 6b): projection magnitude covaried with ICI (Spearman’s rank correlation, for Wa, rho=0.17, p<0.001; for Sa, rho=0.21, p<0.05), whereas component 1 did not show a consistent association (for Wa, rho=0.07, p=0.07; for Sa, rho=0.06, p=0.49). Because component 2 projections reversed sign around partner-call offset (Fig. 5), the same absolute projection magnitude could reflect different dominant propagation directions before versus after call termination. Thus, the ICI relationship may be better understood as reflecting either (1) phase-gradient strength within each epoch-specific direction, or (2) the balance between the two opposite propagation directions captured by component 2. Regardless of mechanism, the significant correlation between component 2 and ICI supports the interpretation that this wave indexes a decision-related variable governing when the subject initiates a response. In contrast, correlations with vocal acoustic features (fundamental frequency f0 and duration) were not consistent across monkeys (Spearman’s rank correlation, for Wa, f0: rho = 0.10, p < 0.05, duration: rho = -0.11, p < 0.01; for Sa, f0: rho = 0.10, p = 0.26, duration: rho = 0.13, p = 0.14), suggesting that the traveling-wave components primarily relate to response timing rather than encoding call acoustics.

We next asked whether traveling-wave dynamics relate to vocalization-induced suppression (“vocal suppression”), a well-established reduction in auditory-cortical spiking that begins immediately prior to vocal onset and is thought to reflect sensory prediction of self-generated vocal output^48,63,66,68,70,71^. Consistent with this framework, we observed suppression of high-gamma power, an established proxy for local spiking, in auditory cortex, and additionally detected suppression across broader cortical areas including frontal regions (Fig. 3).

Trial-by-trial, the magnitude of vocal suppression covaried with component 2 (but not component 1), most prominently in frontal and auditory cortices (Fig. 6c). Specifically, stronger component 2 projections corresponding to propagation from parietal/temporal toward frontal cortex were associated with larger high-gamma suppression, suggesting that macro-scale traveling-wave dynamics interact with local cortical computations. These results may support two nonexclusive interpretations: (1) vocal suppression reflects a process coordinated across multiple cortical areas rather than confined to auditory cortex, and/or (2) the traveling wave contributes to distributing prediction- or decision-related signals that modulate local sensory processing during vocalization.

## Discussion

We recorded whole-cortical epidural ECoG in marmosets engaged in antiphonal calling to examine macro-scale neural dynamics that link vocal perception, response selection, and vocal production. Sensory responses to partners’ calls were widespread and prominently expressed in the theta band, and their magnitude depended on communicative context (i.e., whether the subject subsequently produced a call within the interaction window). Context dependence was also evident around subjects’ own vocalizations, where broad theta and high-gamma suppression differentiated interactive from non-interactive productions. Importantly, these distributed modulations were not merely “global state” changes: a traveling-wave decomposition revealed two reproducible propagation modes with distinct timing and context sensitivity, consistent with separable large-scale processes supporting response selection and motor preparation during vocal communication.

### Involvement of the auditory spatial processing pathway, especially the parietal cortex, in the complete naturalistic vocal-interaction cycle

A prominent finding of our study is the engagement of the auditory–parietal pathway, with a significant contribution of the parietal cortex, during naturalistic vocal exchanges. Parietal cortex showed robust sensory responses to partner calls and strong context-dependent modulation associated with the subject’s subsequent behavior, indicating that parietal activity is not simply a downstream reflection of auditory activation but is engaged in computations relevant to interactive communication. This result aligns with evidence that caudal auditory and temporo-parietal regions carry spatially selective auditory signals^82–86^ and that parietal cortex can represent decision variables in action selection^87,88^.

We propose that our antiphonal calling paradigm specifically recruits these parietal computations because it requires subjects to operate under conditions that closely resemble real-world communication demands: calls are externally generated by the partner, behaviorally meaningful, and embedded in a social exchange in which the subject must determine whether to respond and when to do so. Importantly, subjects were visually isolated from their partner, increasing reliance on auditory cues for localization, source segregation, and interaction timing^89^. Under this framework, parietal cortex may act as a hub that integrates auditory input with spatial/contextual information, transforming sensory evidence into a response policy that governs the initiation and timing of antiphonal vocalizations.

More broadly, the parietal involvement we observe highlights the importance of studying the complete communication process, from perception to response selection to production, within naturalistic social exchanges rather than passive listening alone^90,91^. Parietal activation is not consistently prominent in fMRI studies using passive auditory stimulation^92–95^, which may reflect the fact that these paradigms often minimize the need for spatial inference, turn-taking, or action selection. By contrast, interactive calling imposes immediate behavioral relevance and demands sensorimotor integration, conditions under which parietal contributions are expected to emerge. Thus, our results suggest that a whole-cortex approach applied to interactive vocal behavior can reveal network elements, such as parietal cortex, that may be underestimated when communication is decomposed into isolated components^96^. An important future direction will be to test this interpretation directly by manipulating the social and spatial demands of the interaction: e.g., allowing visual contact with the partner, reducing spatial uncertainty, or using controlled virtual acoustics.

### Traveling-wave dynamics as a structured form of macro-scale coordination

Beyond regional modulations, our analyses demonstrate that vocal interactions are accompanied by structured macro-scale dynamics that are well described as traveling waves, organized phase-gradient patterns spanning the cortex^97–111^. Traveling-wave phenomena have been reported across measurement modalities and species and have been linked to perception and behavior^97–110,112–114^. For example, spontaneous traveling waves in marmoset middle temporal cortex facilitate visual target detection^74^. Similarly, human traveling wave predicted behavioral task performance^75,76^. Here, traveling-wave components carried behaviorally relevant information: their trial-by-trial expression predicted whether and when subjects responded and related to vocalization-induced suppression during production (Figure 5,6). These findings support the view that traveling waves can serve as an efficient mechanism for coordinating distributed computations, linking sensory analysis to decision formation and motor preparation during social communication.

At a mechanistic level, traveling waves are commonly thought to emerge from interactions between local oscillatory generators (shaped by excitatory–inhibitory circuitry) and specific long-range anatomical connectivity that constrains propagation direction and speed, with non-specific connections which follow an exponential decay in connectivity strength as a function of the physical distance between neurons^98,110,115–120^. In this context, theta-band dynamics are well suited to coordinate cross-areal communication^121,122^. The observed relationship between a global traveling-wave component and localized high-gamma suppression further suggests cross-scale coupling, where macro-scale coordination interacts with local population activity linked to spiking (Figure 6c)^75,120,123,124^.

At the same time, our conclusions are based on functional measures inferred from ECoG and phase-gradient structure. Future work combining causal perturbations and circuit-level recordings will be necessary to establish how these waves are generated and whether they are required for controlling interaction timing^74,98,117,125^. Nevertheless, the specificity of wave direction, timing, and context dependence argues against the simple reflection of volume conduction and supports the interpretation that these patterns reflect genuine structured propagation^126,127^.

### Distinct wave components for response selection and motor preparation

The two dominant traveling-wave components differed in their temporal profiles and behavioral relationships. Component 1 exhibited a slower, sustained modulation after partner-call onset and a pre-vocal ramping pattern that peaked one second before vocal onset, consistent with a motor preparatory process approaching a threshold for vocal initiation^128–131^.

Component 2 expressed earlier context dependence during listening, showed direction switching around partner-call offset, and predicted response timing. In contrast to prior findings that emphasized prospective relationships between neural activity and upcoming call acoustics^48^, component 2 showed limited and inconsistent relationships in our dataset. This suggests that, at the macro-scale level captured by whole-cortex traveling waves, component 2 primarily reflects decision- and timing-related variables (e.g., whether and when to respond) rather than encoding the detailed acoustic structure of the forthcoming call^132^. This apparent dissociation could reflect the difference between global propagation dynamics and more spatially localized acoustic representations, as well as differences between magnitude-based and phase-gradient-based analyses.

Both traveling-wave components already differentiated interactive from non-interactive trials during the partner’s call, suggesting that the cortex may enter an interaction-relevant state while sensory evidence is still being acquired. This interpretation is consistent with an early stage of response selection unfolding during listening, rather than a process that begins only after the partner finishes calling. Strikingly, component 2 exhibited a rapid reversal in propagation direction at the partner’s call offset: anterior-to-posterior during the call, but switching to posterior-to-anterior immediately after offset. This pattern is not necessarily counterintuitive, because traveling-wave direction reflects the propagation of a coordinated theta-phase gradient (a spatiotemporal pattern of excitability or gain modulation) rather than mapping one-to-one onto the dominant direction of sensory information or feedforward vs. feedback information flow. In the interactive condition, the partner’s call is behaviorally meaningful and embedded in an imminent action context, such that feedback signals related to emerging response intention could plausibly shape cortical coordination even while the call is ongoing. The offset-locked reversal may therefore reflect a dynamical transition from stimulus-locked, interaction-dependent coordination during listening to a post-offset state that preferentially engages posterior-to-anterior coordination, consistent with forwarding accumulated auditory, spatial, and contextual information to frontal circuits to determine response timing. In parallel, component 1 was stronger in interactive trials and ramped toward vocal onset, consistent with a complementary role in vocal motor preparation once a response has been selected.

Furthermore, in the pre-vocal period, the second component was related to high-gamma vocal suppression, particularly in frontal and auditory cortices. Because vocal suppression is widely interpreted as reflecting predictive signals about self-generated sounds (and the comparison between expected and received feedback)^48,63,65–71,81,133–139^, this association raises the possibility that the decision-related wave contributes to distributing vocal-timing related predictive information to sensory regions, modulating local cortical responses during production^18,48,71,140–146^, possibly through the cross-frequency coupling between macroscale, theta-band traveling wave and localized high-gamma activity as discussed above^75,120,123,124,147,148^. Notably, the pre-vocal association between component 2 and suppression was observed across auditory, parietal, and frontal sites, consistent with a distributed production-ready state rather than a purely local sensory effect.

An alternative interpretation is that these wave components reflect changes in attention or arousal. Although attention likely accompanies social interaction, several observations argue against a purely attention or arousal state-based explanation: context effects were temporally specific rather than sustained, and they manifested in structured propagation patterns rather than uniform gain changes. Future experiments that independently manipulate attention and interaction demands will help disentangle these possibilities.

### Macro-scale dynamics as an integrative mechanism for vocal communication

Finally, whereas we emphasized changes in the macroscale activity pattern during vocal interactions, it should be noted that both waves spontaneously exist: e.g., both waves’ projection values were greater than zero in the baseline period (Fig. 5). Therefore, instead of the emergence of the waves only during vocal communication, spontaneous traveling waves are utilized and modulated during various communicative events. In fact, although context effects were stronger in interactive trials, both wave components were present during non-interactive contexts, suggesting that they may carry general information related to sensory processing and motor output while also gaining access to interaction-specific decision variables. For example, both traveling waves showed sensory responses for partner’s calls even when the subject did not respond to the partner. Also, no clear difference in the traveling wave related to subject’s vocal production between communicative contexts, but build-up (component 1) and decrease (component 2) projection values during pre-vocal and vocal periods implicate vocal-production related shared signals.

Human speech involves distributed processing across temporal and frontal cortices, and directional signaling between these regions has been implicated in speech or vocal planning and production^4,14–18^. Additionally, within-areal local traveling waves facilitate behavioral performances in other cortical areas, including the parietal cortex^74,113^. Our findings extend this framework by showing that such directional interactions can be embedded within broader, whole-cortical traveling-wave dynamics that may integrate perception, decision, and action across a primate cortex during natural communication possibly through gain modulations by interactions of multiple different waves.

## Methods

RIKEN Ethical Committee approved all of the experimental protocols [No. W2020-2-008(4)]. All surgical procedures were conducted using aseptic surgical techniques and with the monkeys kept under general anesthesia as described in Komatsu et al. 2019^72^.

### General

Two adult marmoset monkeys (*Callithrix jacchus*) participated in this study including Wa (male, four years old) and Sa (male, 2 years old). We recorded neural activity from the entire cortical surface using 96-channel epidural ECoG arrays (diameter of each electrode contact: 0.8 mm; Cir-Tech Co. Ltd., Japan) while marmoset monkeys freely produce vocalizations in the laboratory (Figure 2). ECoG arrays were implanted over the left hemisphere for both monkeys. The locations of recording electrodes were identified by merging pre-surgically acquired T2-weighted magnetic resonance images and postoperative computer tomography scans. We estimated the location of each electrode on cortical areas by using a marmoset brain atlas^149^.

### Vocal and neural recording during vocal interactions

We simultaneously recorded neural activity and vocalizations while monkeys freely produced vocalizations in the laboratory. To test interactive vocal behavior, we adopted an antiphonal calling paradigm^57^. This paradigm utilizes marmoset monkeys’ natural vocal behavior when they are visually isolated from others. Specifically, visually-isolated marmosets preferentially produce a specific type of contact call, phee call, and monkeys exchange phees when other partner monkeys respond with phee calls.

In the recording session, we first placed a neural recording “subject” monkey in a small transparent acrylic cage and then brought another “partner” monkey into another cage which is 2.5 m away from the subject cage. We put a curtain to prevent visual contact between them, but allow them to vocally interact freely (Figure 1a). Monkeys engaged in recording sessions for about 90 minutes. We had totally 39 sessions (22 sessions for Wa, 17 sessions for Sa) throughout the whole research.

We recorded vocalizations using handy audio recorders (Zoom H5) placed ∼20cm in front of the subject and partner monkey. Audio data was sampled at 48 kHz. We also recorded videos of up to three cameras to help identify a caller monkey. Offline analysis extracted vocalizations from the recorded signals and classified them into established marmoset call types based on their spectrograms using a semi-automated system. Although most interactive calls were classified as phee calls, we have identified trillphee and trill calls, too, and excluded them for further vocal interaction analysis. ECoG signals were recorded using a wireless neural recording system (CerePlex, Blackrock Microsystems) with a 1 kHz sampling rate. We extracted local field potentials by first subtracting the average activity recorded from all electrodes to reduce muscle potentials and other movement artifacts and then band-pass filtering (1-300 Hz). We removed neural data for some vocalizations due to the loss of wireless recording signals. Since our neurophysiological analyses were restricted to a 12-s peri-event window (−6 to +6 s) aligned to the partner’s call onset, the partner’s call offset, or the subject’s call onset, we defined “signal loss” as data in which the across-channel standard deviation computed in 10-ms bins fell below 30 microvolt within this window.

### Behavioral data analysis

Vocalizations were first detected and extracted from recordings using a semi-automated method based on crossings of a sound amplitude threshold. Vocalizations were then classified into established marmoset call types based on visual inspection of the spectrograms^66,150–152^. We next characterized the acoustic properties and call structure of each vocalization, including duration and fundamental frequency (*f*0), and vocal amplitude. For the *f*0, we first generated call spectrograms to identify a frequency with the maximum power in each time bin and then averaged the frequencies over time. Although the vocal amplitude was calculated as a root mean square amplitude averaged across the entire vocal duration, we noticed the phee calls surpass the dynamic range of recording systems, limiting the reliable measurements, and did not include vocal amplitude for the further data analysis.

To test whether the subject monkey interacted with the partner, we calculated the timing between successive calls, inter-call interval (ICI)^58,153^. Intervals were calculated between the offset time of vocalization and the onset time of the following call. We are particularly interested in inter-call intervals between partners’ preceding calls and subject monkeys’ following calls. To statistically test the ICI distributions from random, we shuffled the timing of vocalizations and generated a null distribution and repeated this procedure for 10000 times to get a chance distribution. We used 95% confidence interval of the chance distribution as a statistical criterion. To estimate the spontaneous vocal production rate, we also calculated the onset-to-onset interval of subject monkeys’ successive calls.

### Neural data analysis

All neural analyses used custom MATLAB and python scripts incorporating functions from the Dr. Kahana lab EEG toolbox (https://memory.psych.upenn.edu/Software). Vocal production–related ECoG activity was visualized with Morlet wavelet time–frequency transforms (6-cycle width; 4–150 Hz). For each event, we computed the wavelet spectrogram and then averaged across vocalizations.

Based on prior reports of pre-vocal suppression and inspection of ECoG signal, we defined: baseline (−3.0 to −2.5 s relative to vocal onset), pre-vocal (−1 to 0 s), and vocal periods (call duration)^48,63,66,70,71^. Pre-vocal and vocal ECoG signals were normalized by dividing with baseline activity. Time–frequency analyses revealed activity related to sensory processing and vocal production in theta plus high-gamma ranges: theta band 5-8 Hz and high-gamma band 50-150 Hz. To quantify these, we band-pass filtered (four-pole Butterworth), applied the Hilbert transform, and normalized amplitudes to baseline.

For theta and high-gamma activity, we first tested whether each site’s neural activity during listening partners’ vocalizations is different depending upon whether subject monkeys vocally respond to them (interactive vs. no interactive) using Wilcoxon rank-sum test (p<0.05). We further quantified the degree of context preference by calculating receiver-operating characteristic (ROC) curves with a permutation test with false discovery rate (FDR) correction (assessing deviation from chance using 95% confidence intervals of 1,000 shuffled data). We assigned ROC>0.5 (<0.5) as greater (smaller) activity in the interactive condition. We further tested the relationship of ROC-based context preference between theta and high-gamma band activity across all recording sites using Spearman’s rank correlation test. Similarly, we also tested whether theta and high-gamma activity depend on communicative context during pre-vocal and vocal periods.

Based on site-by-site modulations by communicative context, we next tested whether and how macro scale cortical activity patters change. Here, we focused on theta-band activity because it is suitable to detect interactions across cortical areas and have been previously used for long-range connectivity analyses^48,121,122^. We first applied a correlation-based functional connectivity analysis^154^. We separately calculated activity relationships across recording sites using Spearman’s rank correlation during partner’s call, pre-vocal, and vocal epochs. The correlation pattern was further classified using a hierarchical clustering and visualized as a dendrogram.

Next, we tested the complex activity propagation patterns on call-by-call basis with fine temporal resolution by extracting macroscale activity propagation patterns. We applied a recently developed method to decomposes the entire-cortical oscillations into directionally independent phase gradient by using the weakly orthogonal conjugate contrast analytical method (WOCCA)^77^. Similar to the principal component analysis, this analysis gives us traveling wave components with explained traveling energy (similar as explained variance in PCA), allowing us to extract the pattern and contribution of each wave). For each wave component, the sign of projection values defines the dominant direction, while their magnitude quantifies the strength of the wave pattern. We compared projection values across communicative context using Wilcoxon rank-sum test during partner’s call, pre-vocal, and vocal periods.

To test the behavioral significance of the traveling waves, we examined the relationships between each wave’s trial-by-trial projection values with behavioral measurements, including vocal production onset and ICI. For vocal onset, we tested whether projection values exhibit a significant peak shortly before vocal production. We randomly selected 120 vocalizations, averaged projection values, and detected a peak time. We repeated this procedure 1500 times and compared with the distribution obtained from temporally shuffled data set (shuffled 1000 times). We tested the statistical significance of the original peak distribution based on the 95% confidence interval of shuffled distribution. For ICI, we calculated Spearman’s rank correlations between ICI and mean projection values from partner’s call offset to 1 s.

Finally, we tested the relationship between traveling waves and local neural activity. More specifically, we focused on the role of traveling waves in suppressive high-gamma activity observed in the pre-vocal period. We calculated Spearman’s rank correlations between projection values and high-gamma activity for each recording site.

Unless otherwise noted, all statistical tests were performed using non-parametric methods, with FDR correction for multiple comparisons if applicable. P-values <0.05 were considered statistically significant.

## Acknowledgement

We woud like to thank Yuri Shinomoto for technical assistance in animal experiments and RIKEN ARD/RRD for marmoset care. We would also like to thank several rotation and internship students in CIBR to support pre-processing and the early part of the data analysis. We thank Drs. Yichao Li and Bo Hong for their scientific, technical advices on the traveling wave analysis. We are grateful for the support provided by the Tsinghua University Non-Human Primate Research Center.

This work was supported by the Beijing Natural Science Foundation IS24043 (J.T.), National Natural Science Foundation of China 32271083 (J.T.), and Japan Science and Technology Agency, PREST JPMJPR22S6 (MK).

## Author contributions

D.Y., M.K., and J.T. designed research; D.Y., M.K., and J.T. performed research; D.Y., X.G., R.T. and J.T. analyzed data; All the authors wrote and edited the paper.

## Declaration of interests

The authors declare that they have no competing interests.

## Supplementary Figure Legends

**Figure S1.**
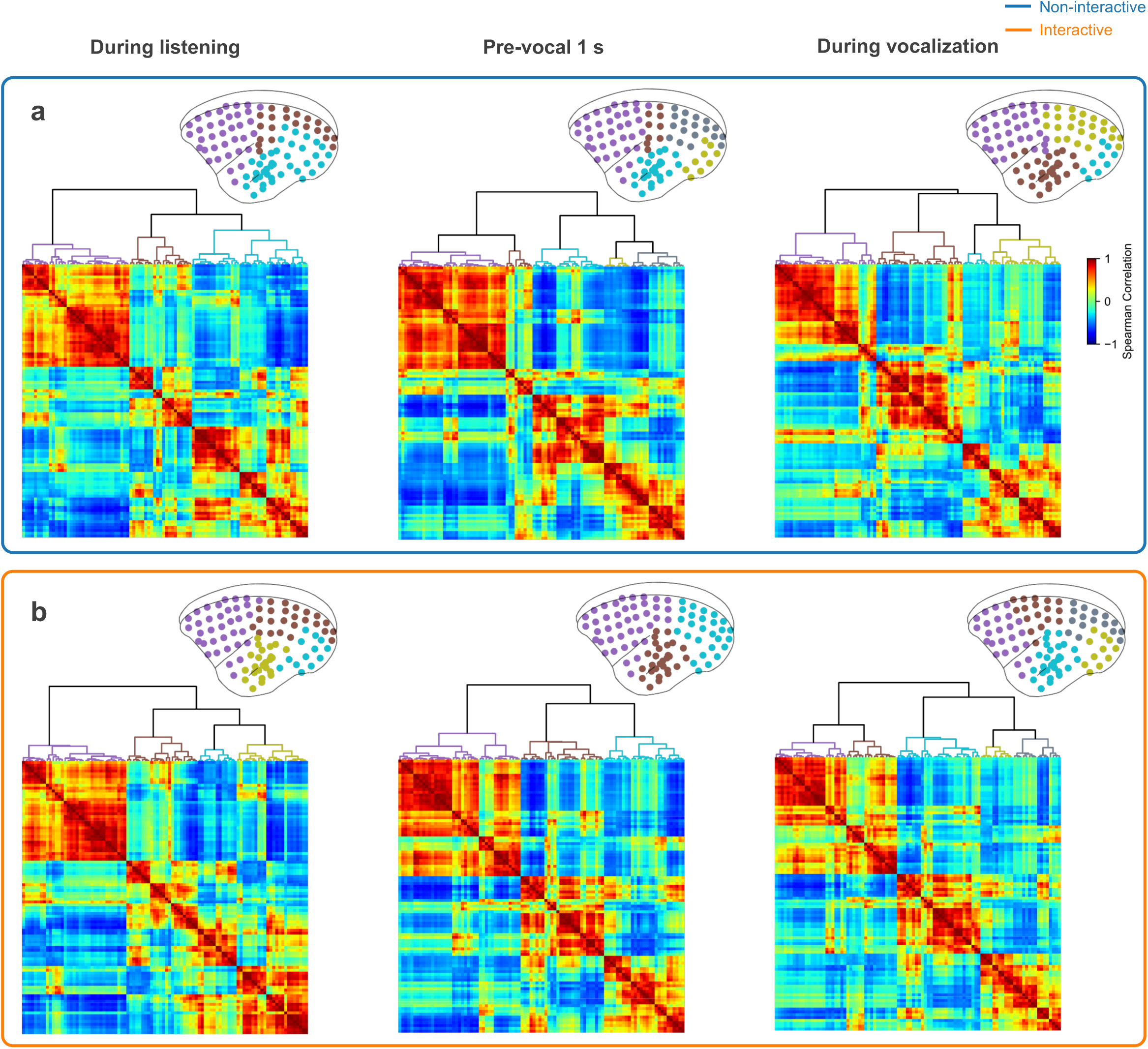
Correlation-based functional connectivity analysis does not differ between interactive and non-interactive conditons. Correlation-based functional connectivity matrices for the non-interactive (a) and interactive (b) conditions. Spearman’s rank correlation was used. The confusion matrices and corresponding hierarchical-clustering dendrograms are shown during the partner’s call (left), pre-vocal (1 s before subject vocal onset; middle), and vocalization (right) periods. Dendrograms were generated by hierarchical clustering of the confusion matrices. The distance threshold was detemined using the elbow method. Clusters with linkage distance below this threshold were grouped and visualized on the cortex using the same color.

**Figure S2.**
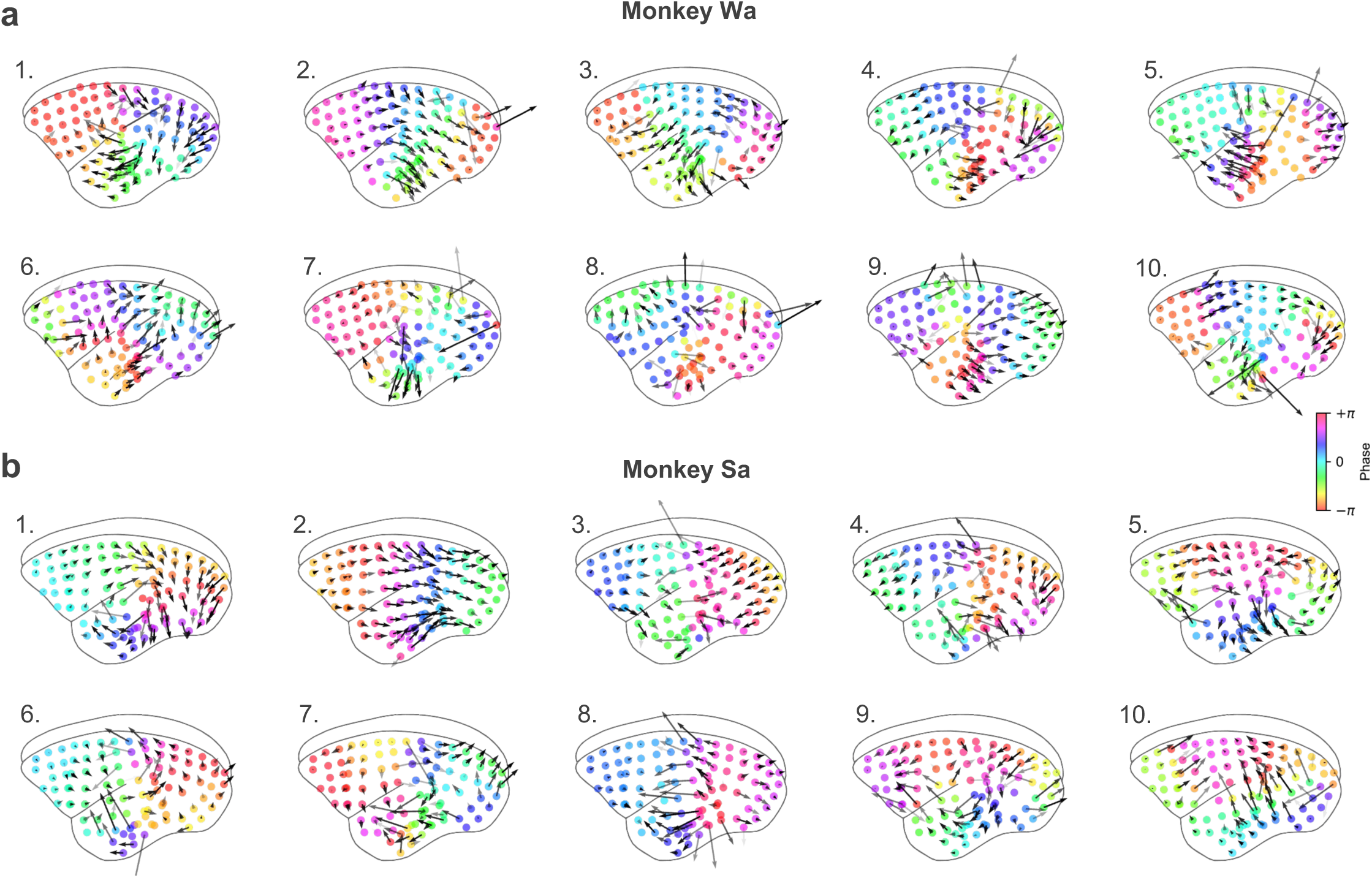
Top 10 WOCCA components. Macroscale theta-phase gradient patterns for the top 10 WOCCA components, ordered by explained traveling energy, for Monkey Wa (a) and Sa (b). The main analyses use the top two components.

**Table S1.**
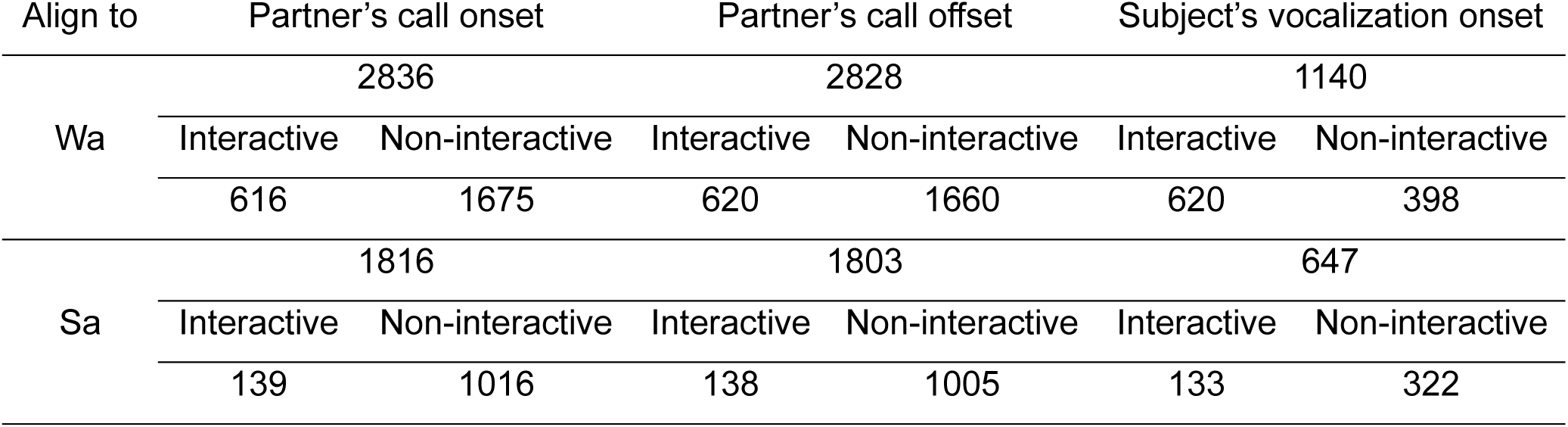
Call numbers used for each epoch.

| Align to | Partner's call onset |  | Partner's call offset |  | Subject's vocalization onset |  |
| --- | --- | --- | --- | --- | --- | --- |
|  | 2836 |  | 2828 |  | 1140 |  |
| Wa | Interactive | Non-interactive | Interactive | Non-interactive | Interactive | Non-interactive |
|  | 616 | 1675 | 620 | 1660 | 620 | 398 |
|  | 1816 |  | 1803 |  | 647 |  |
| Sa | Interactive | Non-interactive | Interactive | Non-interactive | Interactive | Non-interactive |
|  | 139 | 1016 | 138 | 1005 | 133 | 322 |

## Notes

### Competing Interest Statement

The authors have declared no competing interest.

## References

1. Cheney, D.L., and Seyfarth, R.M. (1990). How monkeys see the world (University of Chicago Press).

2. Hauser, M.D. (1997). The Evolution of Communication (MIT Press).

3. Seyfarth, R.M., and Cheney, D.L. (2003). Signalers and receivers in animal communication. Annu Rev Psychol 54, 145–173.

4. Castellucci, G.A., Guenther, F.H., and Long, M.A. (2022). A Theoretical Framework for Human and Nonhuman Vocal Interaction. Annu Rev Neurosci 45, 295–316. 10.1146/annurev-neuro-111020-094807.

5. Banerjee, A., and Vallentin, D. (2022). Convergent behavioral strategies and neural computations during vocal turn-taking across diverse species. Curr Opin Neurobiol 73, 102529. 10.1016/j.conb.2022.102529.

6. Grijseels, D.M., Prendergast, B.J., Gorman, J.C., and Miller, C.T. (2023). The neurobiology of vocal communication in marmosets. Ann N Y Acad Sci 1528, 13–28. 10.1111/nyas.15057.

7. Sacks, H., Schegloff, E., and Jefferson, G. (1974). A simplest systematic for the organization of turn-taking for conversation. Language 50, 696–735. 10.1353/lan.1974.0010.

8. Garrod, S., and Pickering, M.J. (2004). Why is conversation so easy? Trends Cogn Sci 8, 8–11. S136466130300295X [pii].

9. Stivers, T., Enfield, N.J., Brown, P., Englert, C., Hayashi, M., Heinemann, T., Hoymann, G., Rossano, F., de Ruiter, J.P., Yoon, K.E., and Levinson, S.C. (2009). Universals and cultural variation in turn-taking in conversation. Proc Natl Acad Sci U S A 106, 10587–10592. 10.1073/pnas.0903616106.

10. Pickering, M.J., and Garrod, S. (2013). An integrated theory of language production and comprehension. Behav Brain Sci 36, 329–347. 10.1017/S0140525X12001495.

11. Henry, L., Craig, A.J., Lemasson, A., and Hausberger, M. (2015). Social coordination in animal vocal interactions. Is there any evidence of turn-taking? The starling as an animal model. Front Psychol 6, 1416. 10.3389/fpsyg.2015.01416.

12. Levinson, S.C. (2016). Turn-taking in Human Communication - Origins and Implications for Language Processing. Trends Cogn Sci 20, 6–14. 10.1016/j.tics.2015.10.010.

13. Grijseels, D.M., Fairbank, D.A., and Miller, C.T. (2024). A model of marmoset monkey vocal turn-taking. Proc Biol Sci 291, 20240150. 10.1098/rspb.2024.0150.

14. Castellucci, G.A., Kovach, C.K., Howard, M.A., 3rd, Greenlee, J.D.W., and Long, M.A. (2022). A speech planning network for interactive language use. Nature 602, 117-122. 10.1038/s41586-021-04270-z.

15. Castellucci, G.A., Kovach, C.K., Tabasi, F., Christianson, D., Greenlee, J.D.W., and Long, M.A. (2024). Stimulation of caudal inferior and middle frontal gyri disrupts planning during spoken interaction. Curr Biol 34, 2719–2727 e2715. 10.1016/j.cub.2024.04.080.

16. Castellucci, G.A., MacKay, M., Kovach, C.K., Tabasi, F., Greenlee, J.D.W., and Long, M.A. (2025). Neural activity flows through cortical subnetworks during speech production. bioRxiv. 10.1101/2025.06.20.660783.

17. Long, M.A., Katlowitz, K.A., Svirsky, M.A., Clary, R.C., Byun, T.M., Majaj, N., Oya, H., Howard, M.A., 3rd, and Greenlee, J.D. (2016). Functional Segregation of Cortical Regions Underlying Speech Timing and Articulation. Neuron 89, 1187-1193. 10.1016/j.neuron.2016.01.032.

18. Flinker, A., Korzeniewska, A., Shestyuk, A.Y., Franaszczuk, P.J., Dronkers, N.F., Knight, R.T., and Crone, N.E. (2015). Redefining the role of Broca’s area in speech. Proc Natl Acad Sci U S A 112, 2871–2875. 10.1073/pnas.1414491112.

19. Wang, X. (2000). On cortical coding of vocal communication sounds in primates. PNAS 97, 11843–11849.

20. Rauschecker, J.P., and Scott, S.K. (2009). Maps and streams in the auditory cortex: nonhuman primates illuminate human speech processing. Nat Neurosci 12, 718–724. 10.1038/nn.2331.

21. Romanski, L.M., and Goldman-Rakic, P.S. (2002). An auditory domain in primate prefrontal cortex. Nat Neurosci 5, 15–16.

22. Romanski, L.M., and Averbeck, B.B. (2009). The Primate Cortical Auditory System and Neural Representation of Conspecific Vocalizations. Annu Rev Neurosci 32, 315–346. 10.1146/annurev.neuro.051508.135431.

23. Petkov, C.I., Kayser, C., Steudel, T., Whittingstall, K., Augath, M., and Logothetis, N.K. (2008). A voice region in the monkey brain. Nat Neurosci. 10.1038/nn2043.

24. Perrodin, C., Kayser, C., Logothetis, N.K., and Petkov, C.I. (2011). Voice cells in the primate temporal lobe. Curr Biol 21, 1408–1415. 10.1016/j.cub.2011.07.028.

25. Sadagopan, S., Temiz-Karayol, N.Z., and Voss, H.U. (2015). High-field functional magnetic resonance imaging of vocalization processing in marmosets. Sci Rep 5, 10950. 10.1038/srep10950.

26. Price, C.J. (2012). A review and synthesis of the first 20 years of PET and fMRI studies of heard speech, spoken language and reading. Neuroimage 62, 816–847. 10.1016/j.neuroimage.2012.04.062.

27. Hamilton, L.S., Oganian, Y., Hall, J., and Chang, E.F. (2021). Parallel and distributed encoding of speech across human auditory cortex. Cell 184, 4626–4639 e4613. 10.1016/j.cell.2021.07.019.

28. Mesgarani, N., Cheung, C., Johnson, K., and Chang, E.F. (2014). Phonetic Feature Encoding in Human Superior Temporal Gyrus. Science. 10.1126/science.1245994.

29. Mesgarani, N., and Chang, E.F. (2012). Selective cortical representation of attended speaker in multi-talker speech perception. Nature 485, 233–236. 10.1038/nature11020.

30. Yi, H.G., Leonard, M.K., and Chang, E.F. (2019). The Encoding of Speech Sounds in the Superior Temporal Gyrus. Neuron 102, 1096–1110. 10.1016/j.neuron.2019.04.023.

31. Moses, D.A., Leonard, M.K., Makin, J.G., and Chang, E.F. (2019). Real-time decoding of question-and-answer speech dialogue using human cortical activity. Nat Commun 10, 3096. 10.1038/s41467-019-10994-4.

32. Chang, E.F., Raygor, K.P., and Berger, M.S. (2015). Contemporary model of language organization: an overview for neurosurgeons. J Neurosurg 122, 250–261. 10.3171/2014.10.JNS132647.

33. Dichter, B.K., Breshears, J.D., Leonard, M.K., and Chang, E.F. (2018). The Control of Vocal Pitch in Human Laryngeal Motor Cortex. Cell 174, 21–31 e29. 10.1016/j.cell.2018.05.016.

34. Hickok, G., and Poeppel, D. (2007). The cortical organization of speech processing. Nat Rev Neurosci 8, 393–402.

35. Cooney, C., Folli, R., and Coyle, D. (2018). Neurolinguistics Research Advancing Development of a Direct-Speech Brain-Computer Interface. iScience 8, 103–125. 10.1016/j.isci.2018.09.016.

36. Bouchard, K.E., Mesgarani, N., Johnson, K., and Chang, E.F. (2013). Functional organization of human sensorimotor cortex for speech articulation. Nature 495, 327–332. 10.1038/nature11911.

37. Hage, S.R. (2018). Dual neural network model of speech and language evolution: new insights on flexibility of vocal production systems and involvement of frontal cortex. Current Opinion in Behavioral Sciences 21, 80–87.

38. Silva, A.B., Liu, J.R., Zhao, L., Levy, D.F., Scott, T.L., and Chang, E.F. (2022). A Neurosurgical Functional Dissection of the Middle Precentral Gyrus during Speech Production. J Neurosci 42, 8416–8426. 10.1523/JNEUROSCI.1614-22.2022.

39. Nieder, A., and Mooney, R. (2020). The neurobiology of innate, volitional and learned vocalizations in mammals and birds. Philos Trans R Soc Lond B Biol Sci 375, 20190054. 10.1098/rstb.2019.0054.

40. Jurgens, U. (2009). The neural control of vocalization in mammals: a review. J Voice 23, 1–10. 10.1016/j.jvoice.2007.07.005.

41. Miller, C.T., Thomas, A.W., Nummela, S.U., and de la Mothe, L.A. (2015). Responses of primate frontal cortex neurons during natural vocal communication. J Neurophysiol 114, 1158–1171. 10.1152/jn.01003.2014.

42. Roy, S., Zhao, L., and Wang, X. (2016). Distinct Neural Activities in Premotor Cortex during Natural Vocal Behaviors in a New World Primate, the Common Marmoset (Callithrix jacchus). J Neurosci 36, 12168–12179. 10.1523/JNEUROSCI.1646-16.2016.

43. Zhao, L., and Wang, X. (2023). Frontal cortex activity during the production of diverse social communication calls in marmoset monkeys. Nat Commun 14, 6634. 10.1038/s41467-023-42052-5.

44. Nummela, S.U., Jovanovic, V., de la Mothe, L., and Miller, C.T. (2017). Social Context-Dependent Activity in Marmoset Frontal Cortex Populations during Natural Conversations. J Neurosci 37, 7036–7047. 10.1523/JNEUROSCI.0702-17.2017.

45. Li, J., Aoi, M.C., and Miller, C.T. (2024). Representing the dynamics of natural marmoset vocal behaviors in frontal cortex. Neuron 112, 3542–3550 e3543. 10.1016/j.neuron.2024.08.020.

46. Gavrilov, N., Hage, S.R., and Nieder, A. (2017). Functional Specialization of the Primate Frontal Lobe during Cognitive Control of Vocalizations. Cell Rep 21, 2393–2406. 10.1016/j.celrep.2017.10.107.

47. Hage, S.R., and Nieder, A. (2013). Single neurons in monkey prefrontal cortex encode volitional initiation of vocalizations. Nat Commun 4. 10.1038/Ncomms3409.

48. Tsunada, J., and Eliades, S.J. (2025). Frontal-auditory cortical interactions and sensory prediction during vocal production in marmoset monkeys. Curr Biol 35, 2307–2322 e2303. 10.1016/j.cub.2025.03.077.

49. Leonard, M.K., Gwilliams, L., Sellers, K.K., Chung, J.E., Xu, D., Mischler, G., Mesgarani, N., Welkenhuysen, M., Dutta, B., and Chang, E.F. (2024). Large-scale single-neuron speech sound encoding across the depth of human cortex. Nature 626, 593–602. 10.1038/s41586-023-06839-2.

50. Khanna, A.R., Munoz, W., Kim, Y.J., Kfir, Y., Paulk, A.C., Jamali, M., Cai, J., Mustroph, M.L., Caprara, I., Hardstone, R., et al. (2024). Single-neuronal elements of speech production in humans. Nature 626, 603–610. 10.1038/s41586-023-06982-w.

51. Tsunada, J., and Cohen, Y.E. (2014). Neural mechanisms of auditory categorization: from across brain areas to within local microcircuits. Front Neurosci 8, 161. 10.3389/fnins.2014.00161.

52. Tsunada, J., Lee, J.H., and Cohen, Y.E. (2011). Representation of speech categories in the primate auditory cortex. J Neurophysiol 105, 2634–2646. 10.1152/jn.00037.2011.

53. Tsunada, J., Lee, J.H., and Cohen, Y.E. (2012). Differential representation of auditory categories between cell classes in primate auditory cortex. J Physiol 590, 3129–3139. 10.1113/jphysiol.2012.232892

54. Eliades, S.J., and Miller, C.T. (2017). Marmoset vocal communication: Behavior and neurobiology. Dev Neurobiol 77, 286–299. 10.1002/dneu.22464.

55. Miller, C.T., Freiwald, W.A., Leopold, D.A., Mitchell, J.F., Silva, A.C., and Wang, X. (2016). Marmosets: A Neuroscientific Model of Human Social Behavior. Neuron 90, 219–233. 10.1016/j.neuron.2016.03.018 S0896-6273(16)30007-1 [pii].

56. Eliades, S.J.T., J. (2019). Marmosets in auditory research. In The common marmoset in captivity and biomedical research, (Academic Press), pp. 451–475.

57. Miller, C.T., and Wang, X. (2006). Sensory-motor interactions modulate a primate vocal behavior: antiphonal calling in common marmosets. J Comp Physiol A Neuroethol Sens Neural Behav Physiol 192, 27–38. 10.1007/s00359-005-0043-z.

58. Miller, C.T., Beck, K., Meade, B., and Wang, X. (2009). Antiphonal call timing in marmosets is behaviorally significant: interactive playback experiments. J Comp Physiol A Neuroethol Sens Neural Behav Physiol 195, 783–789. 10.1007/s00359-009-0456-1.

59. Roy, S., Miller, C.T., Gottsch, D., and Wang, X. (2011). Vocal control by the common marmoset in the presence of interfering noise. The Journal of experimental biology 214, 3619–3629. 10.1242/jeb.056101.

60. Zhao, L., Roy, S., and Wang, X. (2019). Rapid modulations of the vocal structure in marmoset monkeys. Hear Res 384, 107811. 10.1016/j.heares.2019.107811.

61. Takahashi, D.Y., Narayanan, D.Z., and Ghazanfar, A.A. (2013). Coupled Oscillator Dynamics of Vocal Turn-Taking in Monkeys. Curr Biol 23, 2162–2168. 10.1016/j.cub.2013.09.005.

62. Liao, D.A., Zhang, Y.S., Cai, L.X., and Ghazanfar, A.A. (2018). Internal states and extrinsic factors both determine monkey vocal production. Proc Natl Acad Sci U S A 115, 3978–3983. 10.1073/pnas.1722426115 1722426115 [pii].

63. Eliades, S.J., and Tsunada, J. (2018). Auditory cortical activity drives feedback-dependent vocal control in marmosets. Nat Commun 9, 2540. 10.1038/s41467-018-04961-8.

64. Eliades, S.J., and Tsunada, J. (2023). Effects of Cortical Stimulation on Feedback-Dependent Vocal Control in Non-Human Primates. Laryngoscope 133 *Suppl 2*, S1–S10. 10.1002/lary.30175.

65. Eliades, S.J., and Tsunada, J. (2025). Vocal Error Monitoring in the Primate Auditory Cortex. J Neurosci 45. 10.1523/JNEUROSCI.0090-25.2025.

66. Eliades, S.J., and Wang, X. (2003). Sensory-motor interaction in the primate auditory cortex during self-initiated vocalizations. J Neurophysiol 89, 2194–2207.

67. Eliades, S.J., and Wang, X. (2013). Comparison of auditory-vocal interactions across multiple types of vocalizations in marmoset auditory cortex. J Neurophysiol 109, 1638–1657. 10.1152/jn.00698.2012.

68. Eliades, S.J., and Wang, X. (2008). Neural substrates of vocalization feedback monitoring in primate auditory cortex. Nature 453, 1102–1106. 10.1038/nature06910.

69. Eliades, S.J., and Wang, X. (2005). Dynamics of auditory-vocal interaction in monkey auditory cortex. Cereb Cortex 15, 1510–1523. 10.1093/cercor/bhi030.

70. Tsunada, J., and Eliades, S.J. (2020). Dissociation of Unit Activity and Gamma Oscillations during Vocalization in Primate Auditory Cortex. J Neurosci 40, 4158–4171. 10.1523/JNEUROSCI.2749-19.2020.

71. Tsunada, J., Wang, X., and Eliades, S.J. (2024). Multiple processes of vocal sensory-motor interaction in primate auditory cortex. Nat Commun 15, 3093. 10.1038/s41467-024-47510-2.

72. Komatsu, M., Kaneko, T., Okano, H., and Ichinohe, N. (2019). Chronic Implantation of Whole-cortical Electrocorticographic Array in the Common Marmoset. J Vis Exp. 10.3791/58980.

73. Komatsu, M., Sugano, E., Tomita, H., and Fujii, N. (2017). A Chronically Implantable Bidirectional Neural Interface for Non-human Primates. Front Neurosci 11, 514. 10.3389/fnins.2017.00514.

74. Davis, Z.W., Muller, L., Martinez-Trujillo, J., Sejnowski, T., and Reynolds, J.H. (2020). Spontaneous travelling cortical waves gate perception in behaving primates. Nature 587, 432–436. 10.1038/s41586-020-2802-y.

75. Zhang, H., Watrous, A.J., Patel, A., and Jacobs, J. (2018). Theta and Alpha Oscillations Are Traveling Waves in the Human Neocortex. Neuron 98, 1269–1281 e1264. 10.1016/j.neuron.2018.05.019.

76. Mohan, U.R., Zhang, H., Ermentrout, B., and Jacobs, J. (2024). The direction of theta and alpha travelling waves modulates human memory processing. Nat Hum Behav 8, 1124–1135. 10.1038/s41562-024-01838-3.

77. Li, Y., Qu, J., Yi, D., and Hong, B. (2025). Resolving whole-brain alpha traveling waves across brain states. Neuroimage 322, 121538. 10.1016/j.neuroimage.2025.121538.

78. Julien, C. (2006). The enigma of Mayer waves: Facts and models. Cardiovasc Res 70, 12–21. 10.1016/j.cardiores.2005.11.008.

79. Borjon, J.I., Takahashi, D.Y., Cervantes, D.C., and Ghazanfar, A.A. (2016). Arousal dynamics drive vocal production in marmoset monkeys. Journal of Neurophysiology 116, 753–764. 10.1152/jn.00136.2016.

80. Risueno-Segovia, C., Koc, O., Champeroux, P., and Hage, S.R. (2022). Cardiovascular mechanisms underlying vocal behavior in freely moving macaque monkeys. iScience 25, 103688. 10.1016/j.isci.2021.103688.

81. Greenlee, J.D., Jackson, A.W., Chen, F., Larson, C.R., Oya, H., Kawasaki, H., Chen, H., and Howard, M.A., 3rd (2011). Human auditory cortical activation during self-vocalization. PLoS One 6, e14744. 10.1371/journal.pone.0014744.

82. Tian, B., Reser, D., Durham, A., Kustov, A., and Rauschecker, J.P. (2001). Functional specialization in rhesus monkey auditory cortex. Science 292, 290–293.

83. Recanzone, G.H. (2000). Spatial processing in the auditory cortex of the macaque monkey. Proc Natl Acad Sci U S A 97, 11829–11835.

84. Remington, E.D., and Wang, X. (2019). Neural Representations of the Full Spatial Field in Auditory Cortex of Awake Marmoset (Callithrix jacchus). Cereb Cortex 29, 1199–1216. 10.1093/cercor/bhy025.

85. Chen, C., Xu, S., Wang, Y., and Wang, X. (2025). Location-specific neural facilitation in marmoset auditory cortex. Nat Commun 16, 2773. 10.1038/s41467-025-58034-8.

86. Zhou, Y., and Wang, X. (2012). Level dependence of spatial processing in the primate auditory cortex. J Neurophysiol 108, 810–826. 10.1152/jn.00500.2011.

87. Gold, J.I., and Shadlen, M.N. (2007). The neural basis of decision making. Annu Rev Neurosci 30, 535–574. 10.1146/annurev.neuro.29.051605.113038.

88. Roitman, J.D., and Shadlen, M.N. (2002). Response of neurons in the lateral intraparietal area during a combined visual discrimination reaction time task. J Neurosci 22, 9475–9489.

89. Chen, C., Remington, E.D., and Wang, X. (2023). Sound localization acuity of the common marmoset (Callithrix jacchus). Hear Res 430, 108722. 10.1016/j.heares.2023.108722.

90. Jovanovic, V., Fishbein, A.R., de la Mothe, L., Lee, K.F., and Miller, C.T. (2022). Behavioral context affects social signal representations within single primate prefrontal cortex neurons. Neuron 110, 1318–1326 e1314. 10.1016/j.neuron.2022.01.020.

91. Wong, R.K., Selvanayagam, J., Johnston, K., and Everling, S. (2024). Functional specialization and distributed processing across marmoset lateral prefrontal subregions. Cereb Cortex 34. 10.1093/cercor/bhae407.

92. Jafari, A., Dureux, A., Zanini, A., Menon, R.S., Gilbert, K.M., and Everling, S. (2023). A vocalization-processing network in marmosets. Cell Rep 42, 112526. 10.1016/j.celrep.2023.112526.

93. Dureux, A., Zanini, A., and Everling, S. (2024). Mapping of facial and vocal processing in common marmosets with ultra-high field fMRI. Commun Biol 7, 317. 10.1038/s42003-024-06002-1.

94. Dureux, A., Zanini, A., Trapeau, R., Belin, P., and Everling, S. (2025). Functional organization of voice patches in marmosets and cross-species comparisons with macaques and humans. Curr Biol 35, 3869–3882 e3864. 10.1016/j.cub.2025.07.008.

95. Jafari, A., Dureux, A., Zanini, A., Menon, R.S., Gilbert, K.M., and Everling, S. (2025). Unique Cortical and Subcortical Activation Patterns for Different Conspecific Calls in Marmosets. J Neurosci 45. 10.1523/JNEUROSCI.0670-24.2024.

96. Bennur, S., Tsunada, J., Cohen, Y.E., and Liu, R.C. (2013). Understanding the neurophysiological basis of auditory abilities for social communication: A perspective on the value of ethological paradigms. Hear Res 305, 3–9. 10.1016/j.heares.2013.08.008.

97. Adrian, E.D., and Matthews, B.H. (1934). The interpretation of potential waves in the cortex. J Physiol 81, 440–471. 10.1113/jphysiol.1934.sp003147.

98. Leao, A.A.P. (1944). Spreading depression of activity in the cerebral cortex. J Neurophysiol 7, 359–390.

99. Goldman, S., Santelmann, W.F., Jr., Vivian, W.E., and Goldman, D. (1949). Traveling Waves in the Brain. Science 109, 524. 10.1126/science.109.2838.524.

100. Hughes, J.R. (1995). The phenomenon of travelling waves: a review. Clin Electroencephalogr 26, 1–6. 10.1177/155005949502600103.

101. Massimini, M., Huber, R., Ferrarelli, F., Hill, S., and Tononi, G. (2004). The sleep slow oscillation as a traveling wave. J Neurosci 24, 6862–6870. 10.1523/JNEUROSCI.1318-04.2004.

102. Rubino, D., Robbins, K.A., and Hatsopoulos, N.G. (2006). Propagating waves mediate information transfer in the motor cortex. Nat Neurosci 9, 1549–1557. 10.1038/nn1802.

103. Wu, J.Y., Xiaoying, H., and Chuan, Z. (2008). Propagating waves of activity in the neocortex: what they are, what they do. Neuroscientist 14, 487–502. 10.1177/1073858408317066.

104. Takahashi, K., Saleh, M., Penn, R.D., and Hatsopoulos, N.G. (2011). Propagating waves in human motor cortex. Front Hum Neurosci 5, 40. 10.3389/fnhum.2011.00040.

105. Sato, T.K., Nauhaus, I., and Carandini, M. (2012). Traveling waves in visual cortex. Neuron 75, 218–229. 10.1016/j.neuron.2012.06.029.

106. Muller, L., and Destexhe, A. (2012). Propagating waves in thalamus, cortex and the thalamocortical system: Experiments and models. J Physiol Paris 106, 222–238. 10.1016/j.jphysparis.2012.06.005.

107. Muller, L., Chavane, F., Reynolds, J., and Sejnowski, T.J. (2018). Cortical travelling waves: mechanisms and computational principles. Nat Rev Neurosci 19, 255–268. 10.1038/nrn.2018.20.

108. Das, A., Myers, J., Mathura, R., Shofty, B., Metzger, B.A., Bijanki, K., Wu, C., Jacobs, J., and Sheth, S.A. (2022). Spontaneous neuronal oscillations in the human insula are hierarchically organized traveling waves. Elife 11. 10.7554/eLife.76702.

109. Kaneko, T., Komatsu, M., Yamamori, T., Ichinohe, N., and Okano, H. (2022). Cortical neural dynamics unveil the rhythm of natural visual behavior in marmosets. Commun Biol 5, 108. 10.1038/s42003-022-03052-1.

110. Cruddas, J., Pang, J.C., and Fornito, A. (2026). Cortical traveling waves in time and space: Physics, physiology, and psychology. Neuron. 10.1016/j.neuron.2025.12.019.

111. Ermentrout, G.B., and Kleinfeld, D. (2001). Traveling electrical waves in cortex: insights from phase dynamics and speculation on a computational role. Neuron 29, 33–44. 10.1016/s0896-6273(01)00178-7.

112. Zabeh, E., Foley, N.C., Jacobs, J., and Gottlieb, J.P. (2023). Beta traveling waves in monkey frontal and parietal areas encode recent reward history. Nat Commun 14, 5428. 10.1038/s41467-023-41125-9.

113. Benigno, G.B., Budzinski, R.C., Davis, Z.W., Reynolds, J.H., and Muller, L. (2023). Waves traveling over a map of visual space can ignite short-term predictions of sensory input. Nat Commun 14, 3409. 10.1038/s41467-023-39076-2.

114. Zanos, T.P., Mineault, P.J., Nasiotis, K.T., Guitton, D., and Pack, C.C. (2015). A sensorimotor role for traveling waves in primate visual cortex. Neuron 85, 615–627. 10.1016/j.neuron.2014.12.043.

115. Beurle, R.L. (1956). Properties of a mass of cells capable of regenerating pulses. Philosophical Transactions of the Royal Society London B 240, 55–94.

116. Davis, Z.W., Benigno, G.B., Fletterman, C., Desbordes, T., Steward, C., Sejnowski, T.J., J, H.R., and Muller, L. (2021). Spontaneous traveling waves naturally emerge from horizontal fiber time delays and travel through locally asynchronous-irregular states. Nat Commun 12, 6057. 10.1038/s41467-021-26175-1.

117. Stroh, A., Adelsberger, H., Groh, A., Ruhlmann, C., Fischer, S., Schierloh, A., Deisseroth, K., and Konnerth, A. (2013). Making waves: initiation and propagation of corticothalamic Ca2+ waves in vivo. Neuron 77, 1136–1150. 10.1016/j.neuron.2013.01.031.

118. Canolty, R.T., Edwards, E., Dalal, S.S., Soltani, M., Nagarajan, S.S., Kirsch, H.E., Berger, M.S., Barbaro, N.M., and Knight, R.T. (2006). High power gamma power is phase-locked to theta oscillations in human neocortex. Science 313, 1626–1628.

119. Canolty, R.T., and Knight, R.T. (2010). The functional role of cross-frequency coupling. Trends Cogn Sci 14, 506–515.

120. Bahramisharif, A., van Gerven, M.A., Aarnoutse, E.J., Mercier, M.R., Schwartz, T.H., Foxe, J.J., Ramsey, N.F., and Jensen, O. (2013). Propagating neocortical gamma bursts are coordinated by traveling alpha waves. J Neurosci 33, 18849–18854. 10.1523/JNEUROSCI.2455-13.2013.

121. Buzsaki, G., and Draguhn, A. (2004). Neuronal oscillations in cortical networks. Science 304, 1926–1929. 10.1126/science.1099745.

122. Buzsaki, G. (2006). Rhythms of the Brain (Oxford University Press).

123. Lakatos, P., Shah, A.S., Knuth, K.H., Ulbert, I., Karmos, G., and Schroeder, C.E. (2005). An oscillatory hierarchy controlling neuronal excitability and stimulus processing in the auditory cortex. J Neurophysiol 94, 1904–1911.

124. Hernandez-Perez, J.J., Cooper, K.W., and Newman, E.L. (2020). Medial entorhinal cortex activates in a traveling wave in the rat. Elife 9. 10.7554/eLife.52289.

125. Aggarwal, A., Brennan, C., Luo, J., Chung, H., Contreras, D., Kelz, M.B., and Proekt, A. (2022). Visual evoked feedforward-feedback traveling waves organize neural activity across the cortical hierarchy in mice. Nat Commun 13, 4754. 10.1038/s41467-022-32378-x.

126. Kajikawa, Y., and Schroeder, C.E. (2011). How local is the local field potential? Neuron 72, 847–858. 10.1016/j.neuron.2011.09.029.

127. Orczyk, J.J., Barczak, A., Costa-Faidella, J., and Kajikawa, Y. (2021). Cross Laminar Traveling Components of Field Potentials due to Volume Conduction of Non-Traveling Neuronal Activity in Macaque Sensory Cortices. J Neurosci 41, 7578–7590. 10.1523/JNEUROSCI.3225-20.2021.

128. Hanes, D.P., and Schall, J.D. (1996). Neural control of voluntary movement initiation. Science 274, 427–430.

129. Schurger, A., Sitt, J.D., and Dehaene, S. (2012). An accumulator model for spontaneous neural activity prior to self-initiated movement. Proc Natl Acad Sci U S A 109, E2904–2913. 10.1073/pnas.1210467109.

130. Balasubramanian, K., Papadourakis, V., Liang, W., Takahashi, K., Best, M.D., Suminski, A.J., and Hatsopoulos, N.G. (2020). Propagating Motor Cortical Dynamics Facilitate Movement Initiation. Neuron 106, 526–536 e524. 10.1016/j.neuron.2020.02.011.

131. Balasubramanian, K., Arce-McShane, F.I., Dekleva, B.M., Collinger, J.L., and Hatsopoulos, N.G. (2023). Propagating motor cortical patterns of excitability are ubiquitous across human and non-human primate movement initiation. iScience 26, 106518. 10.1016/j.isci.2023.106518.

132. Liang, W., Balasubramanian, K., Papadourakis, V., and Hatsopoulos, N.G. (2023). Propagating spatiotemporal activity patterns across macaque motor cortex carry kinematic information. Proc Natl Acad Sci U S A 120, e2212227120. 10.1073/pnas.2212227120.

133. Muller-Preuss, P., and Ploog, D. (1981). Inhibition of auditory cortical neurons during phonation. Brain Res 215, 61–76.

134. Creutzfeldt, O., Ojemann, G., and Lettich, E. (1989). Neuronal activity in the human lateral temporal lobe. II. Responses to the subjects own voice. Exp Brain Res 77, 476–489. 10.1007/BF00249601.

135. Numminen, J., Salmelin, R., and Hari, R. (1999). Subject’s own speech reduces reactivity of the human auditory cortex. Neurosci Lett 265, 119–122. 10.1016/s0304-3940(99)00218-9.

136. Houde, J.F., Nagarajan, S.S., Sekihara, K., and Merzenich, M.M. (2002). Modulation of the auditory cortex during speech: an MEG study. J Cogn Neurosci 14, 1125–1138. 10.1162/089892902760807140.

137. Greenlee, J.D., Behroozmand, R., Larson, C.R., Jackson, A.W., Chen, F., Hansen, D.R., Oya, H., Kawasaki, H., and Howard, M.A., 3rd (2013). Sensory-motor interactions for vocal pitch monitoring in non-primary human auditory cortex. PLoS One 8, e60783. 10.1371/journal.pone.0060783.

138. Eliades, S.J., and Wang, X. (2019). Corollary Discharge Mechanisms During Vocal Production in Marmoset Monkeys. Biol Psychiatry Cogn Neurosci Neuroimaging 4, 805–812. 10.1016/j.bpsc.2019.06.008.

139. Chang, E.F., Niziolek, C.A., Knight, R.T., Nagarajan, S.S., and Houde, J.F. (2013). Human cortical sensorimotor network underlying feedback control of vocal pitch. Proc Natl Acad Sci U S A 110, 2653–2658. 10.1073/pnas.1216827110.

140. Khalilian-Gourtani, A., Wang, R., Chen, X., Yu, L., Dugan, P., Friedman, D., Doyle, W., Devinsky, O., Wang, Y., and Flinker, A. (2024). A corollary discharge circuit in human speech. Proc Natl Acad Sci U S A 121, e2404121121. 10.1073/pnas.2404121121.

141. Kingyon, J., Behroozmand, R., Kelley, R., Oya, H., Kawasaki, H., Narayanan, N.S., and Greenlee, J.D. (2015). High-gamma band fronto-temporal coherence as a measure of functional connectivity in speech motor control. Neuroscience 305, 15–25. 10.1016/j.neuroscience.2015.07.069.

142. Behroozmand, R., Shebek, R., Hansen, D.R., Oya, H., Robin, D.A., Howard, M.A., 3rd, and Greenlee, J.D. (2015). Sensory-motor networks involved in speech production and motor control: an fMRI study. Neuroimage 109, 418-428. 10.1016/j.neuroimage.2015.01.040.

143. Behroozmand, R., Oya, H., Nourski, K.V., Kawasaki, H., Larson, C.R., Brugge, J.F., Howard, M.A., 3rd, and Greenlee, J.D. (2016). Neural Correlates of Vocal Production and Motor Control in Human Heschl’s Gyrus. J Neurosci 36, 2302-2315. 10.1523/JNEUROSCI.3305-14.2016.

144. Wang, R., Chen, X., Khalilian-Gourtani, A., Yu, L., Dugan, P., Friedman, D., Doyle, W., Devinsky, O., Wang, Y., and Flinker, A. (2023). Distributed feedforward and feedback cortical processing supports human speech production. Proc Natl Acad Sci U S A 120, e2300255120. 10.1073/pnas.2300255120.

145. Schneider, D.M., Nelson, A., and Mooney, R. (2014). A synaptic and circuit basis for corollary discharge in the auditory cortex. Nature 513, 189–194. 10.1038/nature13724.

146. Schneider, D.M., Sundararajan, J., and Mooney, R. (2018). A cortical filter that learns to suppress the acoustic consequences of movement. Nature 561, 391–395. 10.1038/s41586-018-0520-5.

147. Fries, P. (2015). Rhythms for Cognition: Communication through Coherence. Neuron 88, 220–235. 10.1016/j.neuron.2015.09.034.

148. Fries, P. (2005). A mechanism for cognitive dynamics: neuronal communication through neuronal coherence. Trends Cogn Sci 9, 474–480. 10.1016/j.tics.2005.08.011.

149. Woodward, A., Hashikawa, T., Maeda, M., Kaneko, T., Hikishima, K., Iriki, A., Okano, H., and Yamaguchi, Y. (2018). The Brain/MINDS 3D digital marmoset brain atlas. Sci Data 5, 180009. 10.1038/sdata.2018.9.

150. Epple, G. (1968). Comparative Studies on Vocalization in Marmoset Monkeys (Hapalidae). Folia Primatol 8, 1-&. Doi 10.1159/000155129.

151. Bezerra, B.M., and Souto, A. (2008). Structure and usage of the vocal repertoire of Callithrix jacchus. Int J Primatol 29, 671–701. 10.1007/s10764-008-9250-0.

152. Agamaite, J.A., Chang, C.J., Osmanski, M.S., and Wang, X.Q. (2015). A quantitative acoustic analysis of the vocal repertoire of the common marmoset (Callithrix jacchus). J Acoust Soc Am 138, 2906–2928. 10.1121/1.4934268.

153. Miller, C.T., and Hauser, M.D. (2004). Multiple acoustic features underlie vocal signal recognition in tamarins: antiphonal calling experiments. J Comp Physiol A Neuroethol Sens Neural Behav Physiol 190, 7–19.

154. Friston, K.J., Frith, C.D., Liddle, P.F., and Frackowiak, R.S. (1993). Functional connectivity: the principal-component analysis of large (PET) data sets. J Cereb Blood Flow Metab 13, 5–14. 10.1038/jcbfm.1993.4.

